# Optically Activated, Customizable, Excitable Cells - A Kuhl Platform for Evolving Next Gen Biosensors

**DOI:** 10.1101/2020.02.03.932426

**Authors:** Merrilee Thomas, Thom Hughes

## Abstract

Genetically encoded fluorescent biosensors are powerful tools for studying complex signaling in the nervous system, and now both Ca^2+^ and voltage sensors are available to study the signaling behavior of entire neural circuits. There is a pressing need for improved sensors to properly interrogate these systems. Improving them is challenging because testing them involves low throughput, labor-intensive processes. Our goal was to create a live cell system in HEK293 cells that use a simple, reproducible, optogenetic process for testing prototypes of genetically encoded biosensors.

In this live cell system, blue light activates an adenylyl cyclase enzyme (bPAC) that increases intracellular cAMP [1]. In turn, the cAMP opens a cAMP gated ion channel (olfactory cyclic nucleotide-gated channel, CNG, or the hyperpolarization-activated cyclic nucleotide-gated channel, HCN2). This produces slow, whole-cell Ca^2+^ transients and voltage changes. To increase the speed of these transients, we added the inwardly rectifying potassium channel Kir2.1, the bacterial voltage-gated sodium channel NAVROSD, and Connexin-43. This is a modular system in which the types of channels, and their relative amounts, can be tuned to produce the cellular behavior that is crucial for screening biosensors. The result is a highly reproducible, high-throughput live cell system that can be used to screen voltage and Ca^2+^ sensors in multiple fluorescent wavelengths simultaneously.

## Introduction

### Why are Biosensors Important?

Genetically encoded, fluorescent biosensors are powerful tools for studying cell signaling in real-time [2,3]. They are minimally invasive, and since they are genetically encoded their expression can be exquisitely targeted to specific cell types and tissues [4–6]. However, the short wavelengths of light needed to image green biosensors can heat and damage the brain, decreasing the duration of imaging [7]. There is a need for better biosensors that emit red [8] or near-infrared [9] light because these longer wavelengths enable investigators to image deeper into thick tissues. Furthermore, a better signal to noise ratio can help increase imaging speeds and increase the field of view. Brighter sensors can be detected with lower expression levels potentially decreasing overexpression which can lead to epileptiform activity [10].

A proven approach to expanding the fluorescent biosensor toolbox involves screening thousands of prototypes, evolving for optimal function [11–14]. Ca^2+^ sensors have been highly optimized in *E. coli*. One study used a microfluidic sorter (*μ*FACs) where it was possible to screen up to 10^6^ colonies per round of evolution in *E. coli* [8][15]. A more recent study optimized sensors by picking *E. coli* expressing sensors with the greatest fluorescence intensity. As a second screen, the proteins were cultured and the lysate was placed into Ca^2+^ and Ca^2+^ free buffers and read using a microplate reader [16]. A drawback to this approach is that a sensor cannot be screened for its kinetics, there are only two steady states.

A different approach to screen activity-based biosensors is in mammalian excitable cells. Screening in primary cultures of differentiated, excitable cells is time consuming, expensive, and low throughput(Storace et al. 2016). Another approach is to create excitable cells de novo using transformed cells lines compatible with high throughput screening. One of the first de novo excitable cells was made by expressing rat brain IIA Na^+^ channel and the Drosophila Shaker K^+^ channel into Chinese hamster ovary cells via vaccinia virus [17]. Additionally, one study stably expressed three ion channels, Kir2.1, Nav1.5, and Cx-43 in HEK293 cells. Using electrophysiology they found that with all three channels expressed fast, uniform action potential propagation could be created. However, these particular cells cannot be used to test fluorescent biosensors, because each exogenous channel was marked with a fluorescent protein spanning the visible spectrum [18].

Current methods to screen voltage sensors, in particular, include using excitable HEK293 cells that combine Kir2.1 and a voltage-gated sodium channel (NAV1.3) with field stimulation to produce rhythmic depolarization and hyperpolarization in HEK cells [12,19]; [20]. Another option is to use a stable cell line that expresses Kir2.1, Nav1.7, and CheRiff, a channelrhodopsin. This can be photoactivated thereby creating a system that produces oscillating membrane voltage [21]. Yet, there are inherent problems with stable cell lines, such as genetic drift and gene silencing. Often stable cell lines will regulate the expression of exogenous genes, and mutations in the exogenous genes will occur after several cell culture passages [22,23]. In addition, the level of expression of each of the channels cannot be easily changed and currently, only wavelengths outside of the rhodopsin’s excitation spectrum can be used for screening.

What if we could screen thousands of variants in mammalian cells for sensitivity, speed, and dynamic range? We hypothesized that we could use similar channels as those used previously to create cells that (1) can be optically stimulated (2) produce action potentials and whole-cell Ca^2+^transients and (3) can be imaged with at least two wavelengths simultaneously. We wanted the system to be automatable, inexpensive, modular, and useful in many different cell types. While it could be incorporated into a stable cell line we opted for a more modular route by using baculovirus [24]. While viral transduction involves more work than a stable cell line, it gives us the ability to optimize each component independently, something that cannot be done in stable cells.

Our goal was to create a screening platform for biosensors that is consistent, reliable, and capable of quickly screening thousands of prototypes.

We first created a minimal cellular system with a light-activated enzyme, we used an adenylyl cyclase, to establish a “Kuhl” cell. Next, we added a cAMP-gated ion channel, either a CNG or HCN channel, to produce “Kuhl-C, and Kuhl-H” cells respectively. We then added additional channels and proteins to tune the responses. To improve the responses we added the Kir2.1 potassium channel, to create Kuhl-CK or Kuhl-HK cells. To speed up the action potentials we added the bacterial sodium channel NavD to design Kuhl-CKNa and Kuhl-HKNa cells. Finally, we found that connexin-43 improved the coordination of the signaling, producing Kuhl-CKNaCx and Kuhl-HKNaCx cells.

## Results

### Establishing an Optical Actuator

How can we activate a biosynthetic cell line using light? To develop an optogenetically controlled screen, we chose to use bPAC (blue photo-activated cyclase), a blue light activated adenylyl cyclase from the soil bacteria *Beggiatoia* [1]. In theory, coupling a light activated enzyme to large conductance channels, should produce larger whole cell currents with less protein expression than the current optogenetic approaches that rely upon channel rhodopsins. The bPAC enzyme is activated with 480 nm light and converts ATP into cyclic adenosine monophosphate (cAMP). We began by examining whether blue light stimulation can produce a measurable change in cAMP levels in bPAC-expressing HEK293 cells as seen with R-cADDis, a red fluorescent cAMP sensor [25].

We transduced HEK293 cells with a BacMam virus that expresses bPAC and R-cADDis (S2 Table). The following day, we stimulated cells with 20 milliseconds of blue light and then collected images continuously with 561 nm excitation that does not activate bPAC (Fig 1A). Upon the blue light stimulation, the R-cADDis red fluorescence increases (Fig 1B). cAMP levels within the cell rise over the time frame of ~100 seconds (Fig 1C). Control cells with no bPAC did not show an increase in red fluorescence. The system response is remarkably reproducible and the same cells can be repeatedly stimulated. Indeed the only limitation is that too much blue light given over a short time will create too much cAMP and saturate the sensor. The slow decrease in fluorescence of R-cADDis is most likely due to the phosphodiesterases present in the cell which work to eliminate cAMP [26].

**Fig 1.**
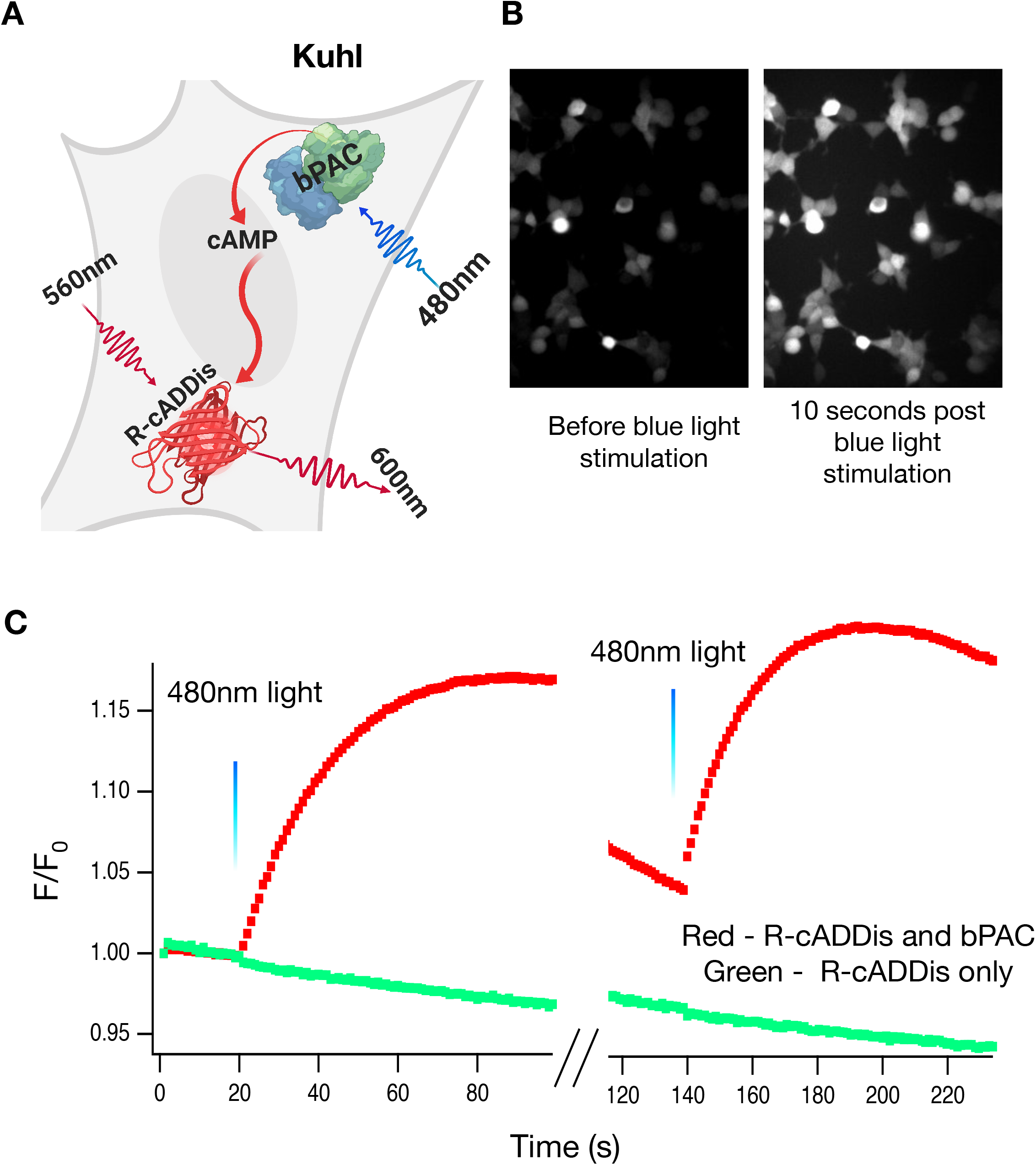
20ms of blue light can repeatedly activate bPAC. A) HEK-293 cells were transduced with bPAC (the actuator) and R-cADDis, the red fluorescence cAMP sensor (S2 Table). B) The cells were illuminated with 20ms of 480nm light. This activation of bPAC increases cAMP levels which increases R-cADDis fluorescence. Brightness and contrast are the same for both images. C) The traces are a discontinuous recording of R-cADDis with 560nm excitation from the same cell (100s on, 20s off, 100s on). 20 ms of blue light at 20s and 140s activates bPAC which produces bursts of cAMP for 100s. This is a stereotyped response that is repeatable with multiple doses of blue light.

### Dose Dependence

To examine the dose-response relationship between blue light and cAMP production, we systematically varied the intensity of the blue light during stimulation while quantifying the R-cADDis fluorescence. HEK293 cells were transduced with bPAC and R-cADDis, and each well in a 96-well plate was stimulated with a 15ms dose of blue light. For each well, LED 480nm light power was increased stepwise from 0 to 70 mW/cm^2^ in increments of five %power using a LED controller. At 100% power, the light intensity at the specimen plane measured 70mW/cm^2^. Figure 2 reveals a dose response in which increasing levels of blue light produces increases in cAMP levels. Lowering the level of bPAC expression by decreasing the total viral load may make it possible to create a system with a greater range of tunability before saturating the sensor with cAMP. The fluorescence change, □F, was used to determine the cAMP level and all points in the dose response are an average of multiple regions of interest (ROIs) within a trial (Fig 2). This is consistent with a similar study that found bPAC had a linear dose response to light when cAMP was recorded [27]

**Fig 2.**
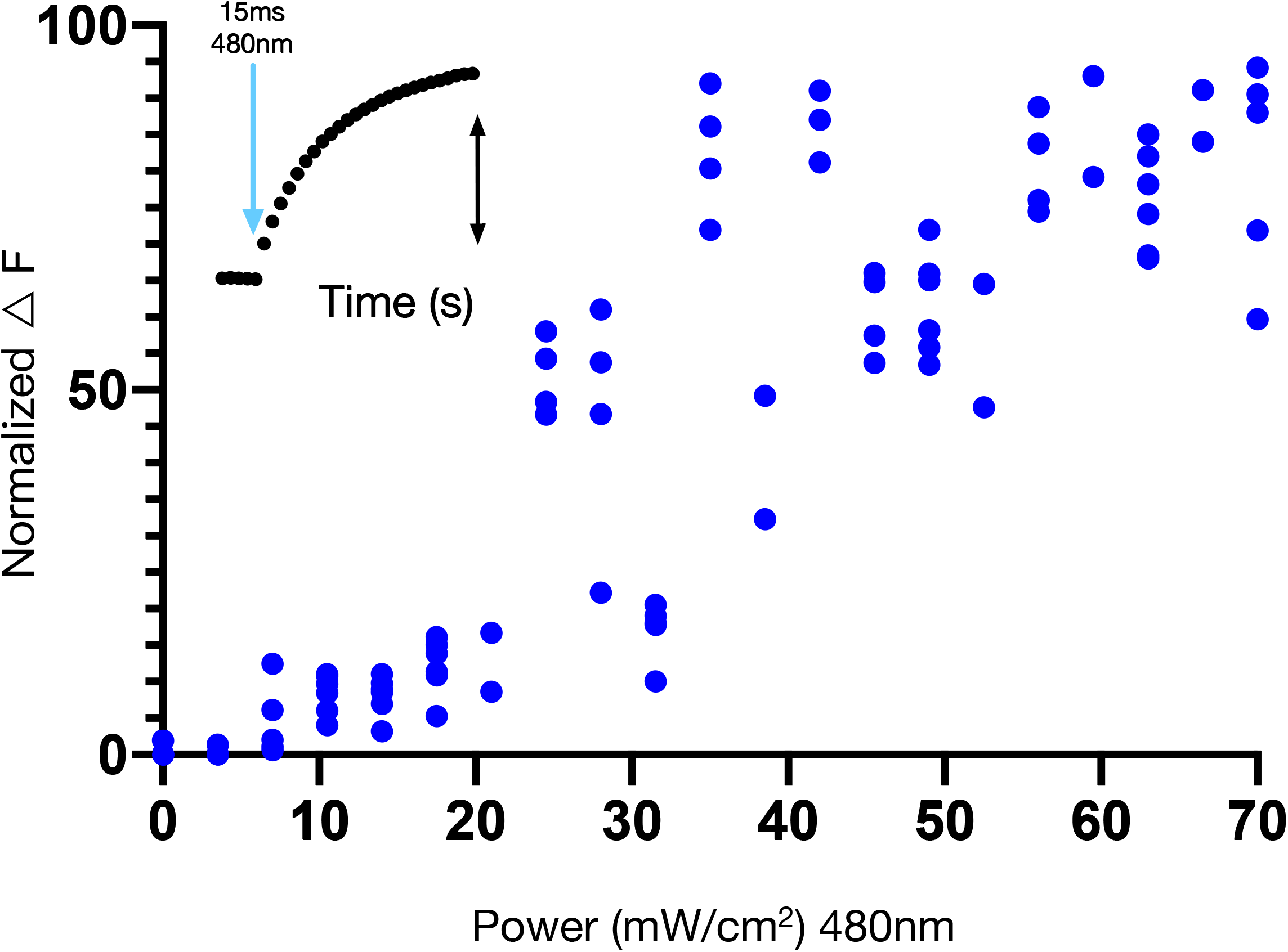
There is a dose/response relationship between the intensity of blue excitation of bPAC and the fluorescence response of R-cADDis. As the blue light intensity increases so does the cAMP production. The □F is a measure between the baseline fluorescence and the maximum of the change in fluorescence (inset). Each data point represents an average of ~100 ROIs within a trial.

### Coupling the Actuator with an Ion Channel

To couple cAMP production with channel activation, we expressed a cyclic nucleotide-gated ion channel (the rat olfactory CNG or human heart HCN2 channel) (S2 Table). The rat olfactory CNG channel is normally gated by cyclic guanosine monophosphate (cGMP) [28]. However, several mutations (561-90/C460W/E583M) can be introduced [29] that renders it sensitive to cAMP. In theory, this mutant channel should couple bPAC activation to a current that depolarizes the cell, creating Kuhl-C cells. To create Kuhl-H cells, we expressed the human HCN2 pacemaker channel which is also cAMP-gated and is hyperpolarization-activated (S2 Table) [30]. Once the cell is hyperpolarized, the HCN2 channel can open when bound to cAMP.

We activated the Kuhl-C or Kuhl-H cells with blue light thereby increasing cAMP. Figure 3 illustrates that 20ms of blue light activation of bPAC, produces an increase in the red fluorescence of R-GECO1, demonstrating a gradual increase in whole-cell Ca^2+^ over several minutes. Opening either the CNG or HCN2 channel increased the red fluorescence of R-GECO1, consistent with increased cytosolic Ca^2+^ levels (Figs 3B and 3F). Interestingly, blue light stimulation also produced bright subcellular flashes of fluorescence in both the Kuhl-C and Kuhl-H cells. The source of this Ca^2+^ is not clear, but one possibility is that increases in cAMP could be causing brief IP_3_ channel opening, or perhaps mitochondrial release [31] (Figs 3C and 3G). The subcellular flashes are small flashes of Ca^2+^ that last ~20 seconds (Figs 3D and 3H). They are far more frequent in cells without NavD but can be seen in all Kuhl cells (S1 Movie). The control cells without bPAC showed no such responses (S1 Fig).

**Fig 3.**
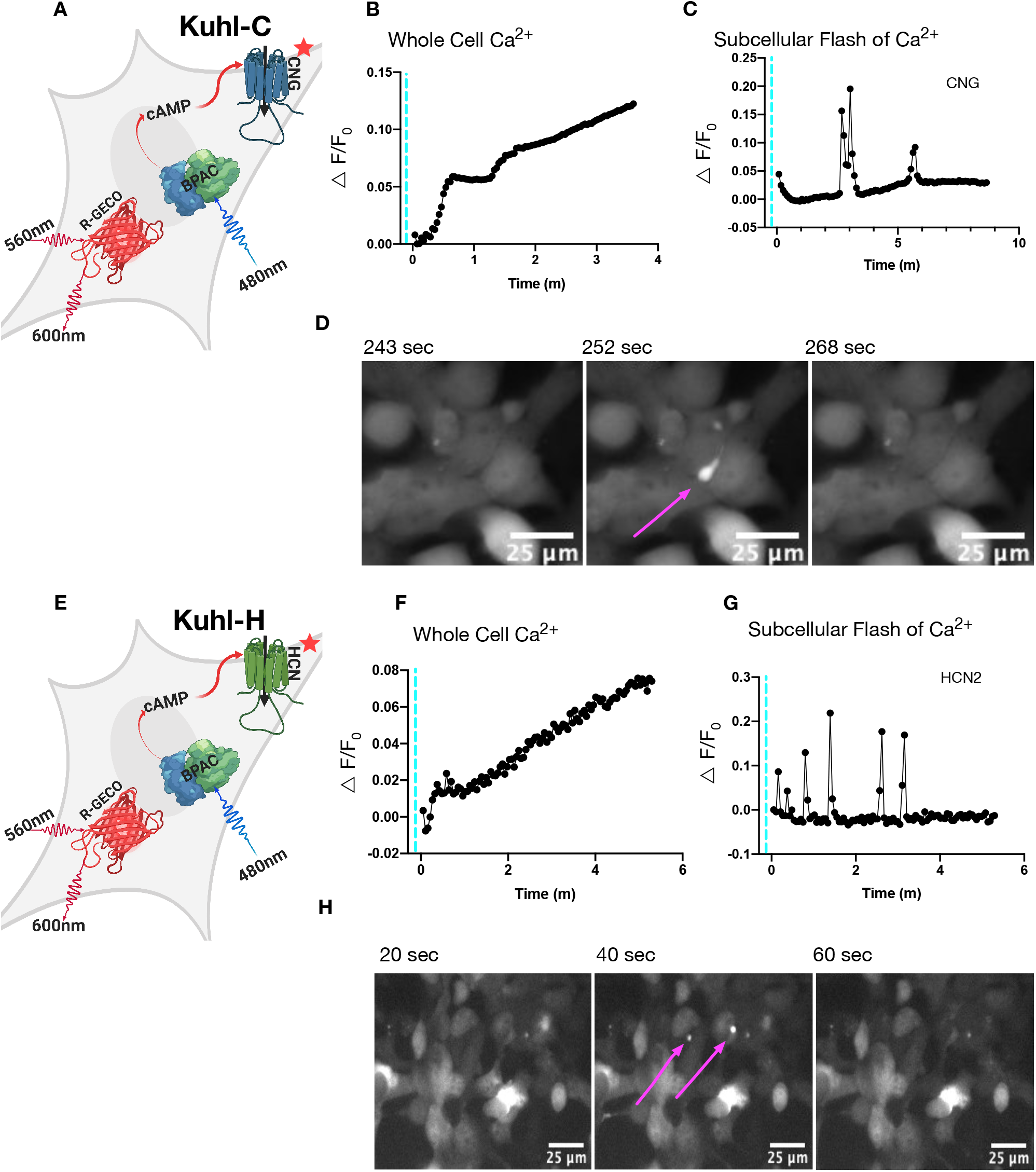
Blue light activation of bPAC slowly increases whole-cell Ca^2+^ when CNG or HCN2 is expressed. There were two types of increases in Ca^2+^ activity: a slow whole-cell Ca^2+^ increase and bright subcellular flashes of Ca^2+^. These can be seen when bPAC and a cyclic nucleotide-gated channel are expressed. A, E) Cartoon depicting all the components in the Kuhl-C or Kuhl-H cell system (S2 Table). Blue light stimulation of bPAC produces cAMP that in turn opens a cAMP gated CNG (delta61–90/C460W/E583M) channel or HCN2 channel. B) The black trace is the R-GECO1 response to 30 ms of blue light activation in cells with both CNG and bPAC, F) or with HCN2 and bPAC. C) The black trace depicts small brief subcellular flashes of Ca^2+^that can be seen in cells with CNG and bPAC, G) or HCN2 and bPAC. D) The localized subcellular flash of Ca^2+^ seen in the Kuhl-C cell system in a series of three frames at 243s, 252s, 268s, and H) in Kuhl-H cells at 20s, 40s, 60s.

### Addition of Kir2.1 Coupled with bPAC’s cAMP Activity Leads to Fluctuating Whole-Cell Ca^2+^ Transients

The bPAC and CNG, or HCN2, expression produced very reliable Ca^2+^ transients, but these were long-lived events with slow kinetics. We hypothesized that introducing a voltage-regulated inwardrectifying potassium channel could speed up, and increase the amplitude, of these transients by creating a greater driving force on Ca^2+^ to enter the cell. Wild type HEK293 cells have a resting membrane potential of only −25 mV, but Kir2.1 expression shifts their resting potential to −70mV [18,32,33]; [34].

We expressed Kir2.1, bPAC, R-GECO1, and a cyclic nucleotide-gated channel in HEK293 cells to produce Kuhl-CK and Kuhl-HK cells (S2 Table). In the Kuhl-C/HK cell system, there is some degree of spontaneous activity before blue light stimulation, but when we activate bPAC with 20ms of blue light there are robust responses in all of the wells. There are Ca^2+^ transients that occur in 85% of the cells, but the activity is largely uncoordinated. Each cell has its own “signature” change in Ca^2+^ levels, as seen in the R-GECO1 fluorescence traces (Fig 4B). Ca^2+^transients can be seen for up to 30 minutes following the blue light stimulus (S2 Fig). The histogram shows the average cumulative number of fluorescence peaks per cell over 340s (Fig 4G). HCN2 produces a slightly greater cumulative average frequency of Ca^2+^ fluorescence peaks than CNG.

**Fig 4.**
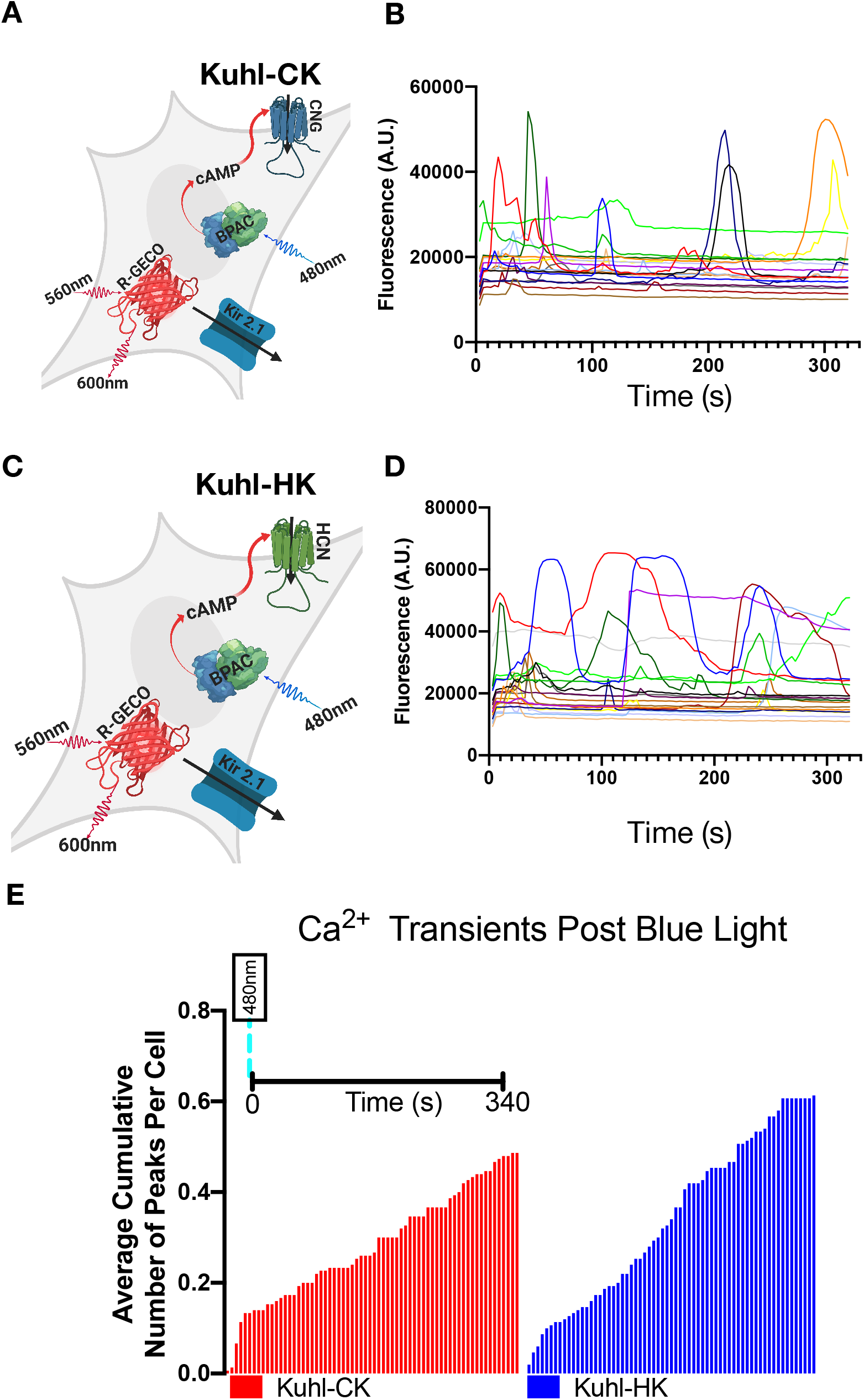
The addition of Kir2.1 leads to fluctuating whole-cell Ca^2+^ transients in the Kuhl-H/CK system. A, C) Cartoon depicting all the components transduced into the experimental Kuhl-H/CK cell system (S2 Table). Blue light stimulates bPAC which will produce cAMP that opens the (CNG or HCN2) channel, and Kir2.1 should hyperpolarize the cells, creating a greater driving force on Ca^2+^ to enter the cell. B) R-GECO1 fluorescence response to 20ms of blue light in Kuhl-CK or D) 20ms of blue light stimulating Kuhl-HK cells. The traces show a subset of the total responses (S2 Fig). G) Average cumulative number of Ca^2+^ peaks in Kuhl-HK and Kuhl-CK system (all peaks detected across 150 cells for each condition). Kuhl-HK cells have a higher average cumulative number of peaks per cell. Each cell has a unique fluctuating Ca^2+^ transient response after the blue light stimulation of bPAC.

### Optimizing the Activity

To quantify the Ca^2+^ imaging data we analyzed it using a custom MATLAB program that harnessed MATLAB’s built-in peak finder (Fig 5A) (methods). We wanted to break down each Ca^2+^ transient into quantifiable components that could be compared across different conditions. We defined a transient as a change in fluorescence that exceeded the standard deviation of the variability in the baseline by a factor of 26. In other words, we only analyzed events where the SNR exceeded 26 [35]. For each fluorescent peak, we quantified the total cumulative peaks (Fig 5b) and the total number of peaks per ROI (Fig 5D). The normalized □F of the peak (amplitude) (Fig 5E), the full width of the peak at half maximum (FWHM) (Fig 5F), and the inter-peak interval (Fig 5G). Also, we examined the total number of Ca^2+^ transients recorded per experiment and recorded the time it took for a Ca^2+^ transient to occur post blue light stimulation.

**Fig 5.**
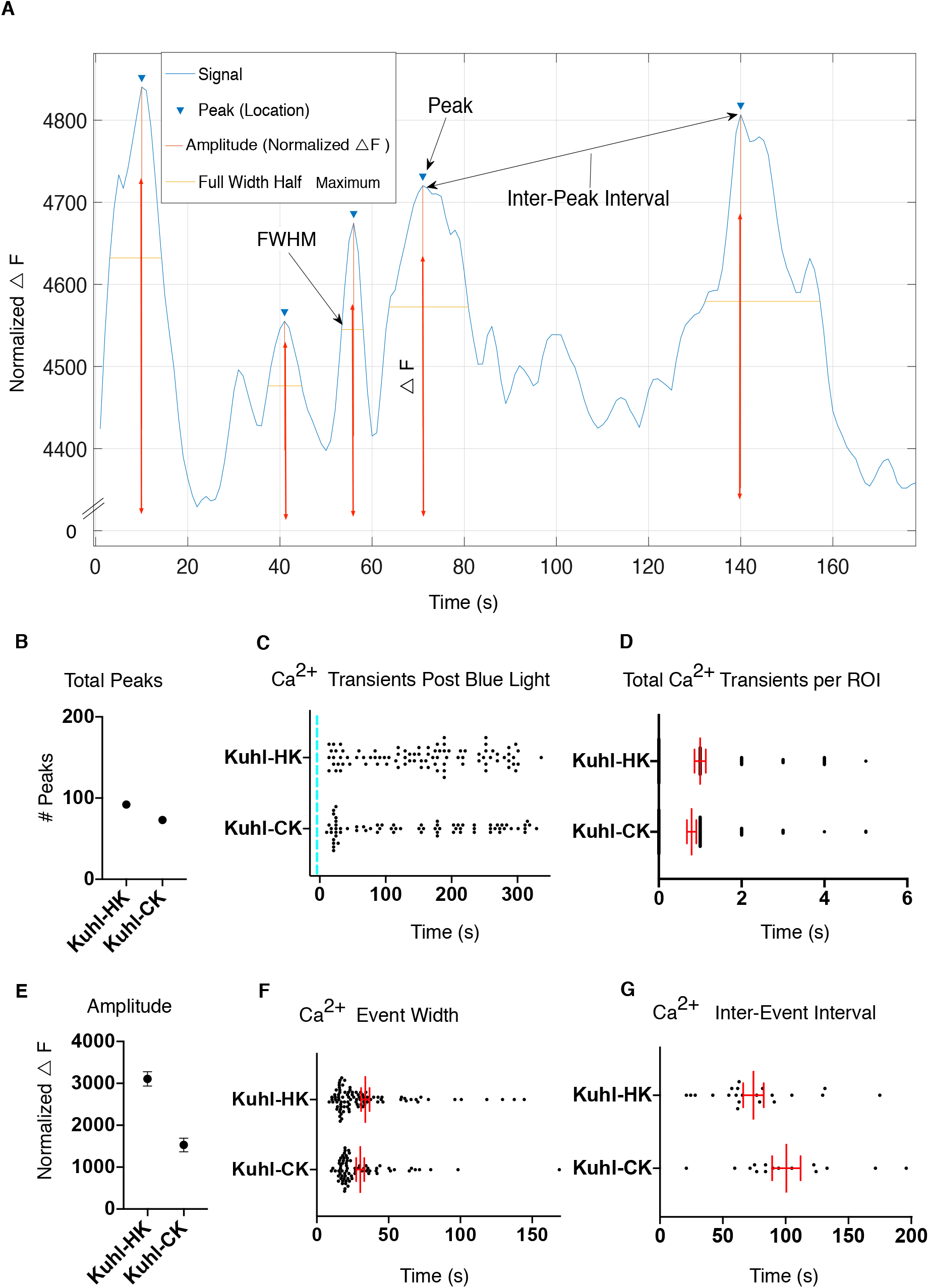
Features of the prolonged Ca^2+^ transients in Kuhl cells shown in Figure 4 differ depending on the presence of CNG or HCN2. A) Diagram of the MATLAB analysis of the Ca^2+^ transients. Peak intensity, the full width of the peak at half maximum (FWHM), and interpeak intervals were analyzed. B) The total Ca^2+^ peaks (*n* = 150 cells per condition) for Kuhl-H/C were analyzed. The Kuhl-HK system had more Ca^2+^ peaks in 350s than the Kuhl-CK system. C) Blue line indicates a point of 20s blue light stimulus. The time of each fluorescence peak was recorded and is indicated by a black dot. D) The number of Ca^2+^transients per cell (ROI) for both HCN2 and CNG. E) The normalized □F is the difference between the baseline fluorescence and the maximum fluorescence of each peak. The mean and S.E.M of the Ca^2+^ transients in Kuhl-H/CK cells. F) Representative duration of elevated R-GECO1 fluorescence overtime per Ca^2+^ transient. FWHM is an estimate of the length of time that the fluorescence stays elevated for each defined peak. G) The Inter-event interval, which is the difference in time between two consecutive peaks. The time between Ca^2+^ transient events per cell. Bars indicate mean and ± S.E.M for 150 cells.

We found that in the Kuhl-HK cell system (*n* = 150 cells) there is a 62% chance that any given cell produces a Ca^2+^ peak (Fig 4G) and when imaged for 340s a 52% chance (methods) that it will produce a second peak within ~70s (Fig 5G) of the initial response. The Kuhl-CK system has a 20% chance that a cell will produce a significant Ca^2+^ peak and out of the cells that were suprathreshold there is a 63% chance that there will be a second response in that cell ~100s (Fig 5G) following.

### Voltage-gated Sodium Channel NavD Creates Faster Ca^2+^ Transients

The bacterial sodium channel, NavRosDg217a (NavD) is characterized by a rapid activation state that depends on the cell depolarizing to −30mV, and a slow inactivation state (hundreds of milliseconds) that starts during the depolarization [32]. Co-expressing the NavD channel with Kir2.1 can potentially create a faster excitable system because the Kir2.1 hyperpolarizes the cell after each spontaneous depolarization caused by the sodium channel. We chose NavD because the mutant is slower than the mammalian sodium channels and in theory, we could create a system with depolarizations such that membrane depolarization events could be captured with slower imaging speeds (10Hz).

A recent paper by Chen et al. found that varying the levels of Kir2.1, and Cav1.3 or Nav1.5 affected the membrane oscillations as recorded by electrophysiology [30]. We hypothesized that different levels of the Kir2.1 and NavD would lead to different corresponding Ca^2+^ responses.

To test this we systematically transduced cells with varying ratios of Kir2.1 and NavD while holding the viral concentrations of bPAC, CNG and R-GECO1 constant. The Kuhl-CKNa cells were transduced (S2 Table) and ~48 hours later the cells were stimulated with 20ms of blue light and images were collected with 561nm excitation. Upon activation of bPAC there were varying responses in each condition that are quantified (S4 Fig).

The Kuhl-CKNa cells with a viral titer of (Kir2.1 1e8):(NavD 5e8) VG (Viral Genomes)/*μ*L had the greatest cumulative peaks(S4a Fig) over 210s. To understand which (Kir2.1):(NavD) ratio had the greatest cumulative number of Ca^2+^ transients divided by the total number of cells (Fig 6b). Using the data from the total number of peaks per ROI (S4d Fig) we ran a Dunnett’s test (S1 Table) to examine if there was a statistical significance between the conditions. (Kir2.1 1e8):(NavD 5e8) VG/*μ*L had a significant increase (\*\*\*\**p* ≤ .0001) in the average cumulative number of peaks per cell (Fig 6b). In the Kuhl-CKNa system, there is a 35% chance that a cell will produce a significant Ca^2+^ peak and out of the cells that were suprathreshold there is a 43% chance of a second peak within 66s of the initial fluorescence peak (S4f Fig).

**Fig 6.**
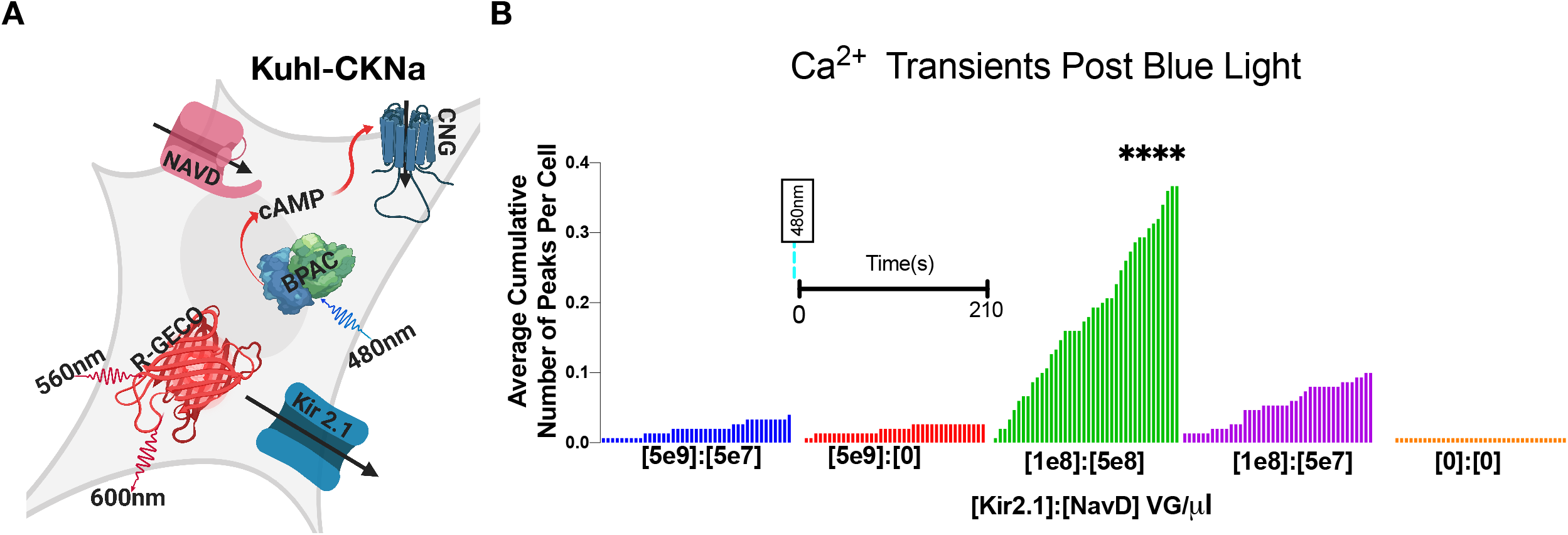
The Kir2.1 and NavD concentration were optimized. NavD was introduced to the Kuhl-CKNa cell system. Introducing NavD created significantly faster Ca^2+^ transients (S3 Fig). A) Cartoon depicting all the components transduced into the experimental Kuhl-CKNa cell (S2 Table). B) Shown is the average cumulative fluorescence peaks per cell across 210s for each indicated viral concentration of Kir2.1 and NavD (*n* = 150 cells per condition). Differences were analyzed using Dunnett’s multiple comparisons test on the Ca^2+^ transient frequency (S4d Fig). (Kir2.1 1e8):(NavD 5e8) VG/*μ*L was significantly greater (\*\*\*\**p* ≤ .0001) than the control (Kir2.1 0):(NavD 0) VG/*μ*l.

Introducing NavD created significantly faster Ca^2+^ transients (S3 Fig). In S3 Figure the differences in the mean duration that Ca^2+^ stayed elevated in the Kuhl-CKNa system with and without NavD are shown. Further optimizing connexin-43 and CNG concentrations significantly decreased the time that Ca^2+^ transients stayed elevated (S3 Fig).

### Further Optimization with Connexin-43 and Kir2.1 and NavD

HEK293 cells have endogenous connexin proteins, connexin-45, which create gap junctions that connect the cytoplasm of the cells [36,37]. However, Kirkton and Bursac [18] found that when a confluent layer of HEK293 cells expressing Kir2.1 and Nav1.5 had action potentials that did not propagate uniformly through the monolayer. When they added Connexin-43 Cx-43 they recorded rapid and robust action potentials. In addition, several other studies have shown that including the Cx-43 protein increases the electrical gap junction coupling of HEK293 cells [18,32,38].

The concentrations for (Kir2.1):(NavD) VG/*μ*L were varied for each condition and the following concentrations were held constant for bPAC, CNG, Cx-43, and R-GECO1 (S2 Table). The Kuhl-CKNaCx cells were transduced (S2 Table) and 48 hours later the cells were stimulated with 20ms of blue light and images were collected with 561nm excitation. Upon activation of bPAC there were varying responses in each condition that are fully quantified (S5 Fig). Interestingly there seems to be a significant effect on the responses in the Kuhl-CKNaCx cells (Fig 7B) depending on the amount of (Kir2.1):(NavD) VG/*μ*L applied.

**Fig 7.**
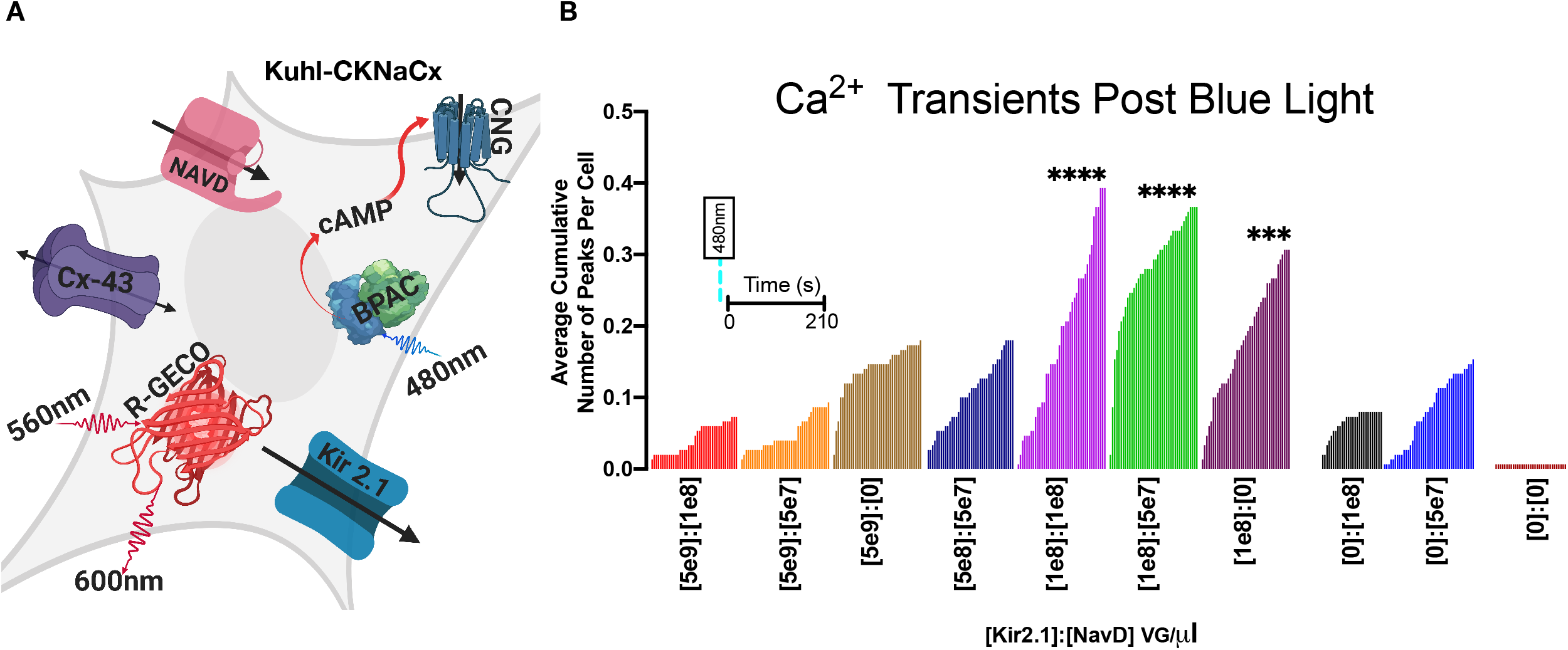
Optimizing Kir2.1 and NavD with Connexin-43 increases the average number of Ca^2+^ transients. Introducing the NavD and Kir2.1 channel is expected to create faster oscillating Ca^2+^ transients in the cell while introducing the Cx-43 will increase coordinated activity between cells. Further optimization of Kir2.1 to NavD is shown. Cx-43 was introduced to the Kuhl-CKNaCx system. A) Cartoon depicting all the components transduced into the experimental Kuhl-CKNaCx cell (S2 Table). B) Shown is the average cumulative fluorescence peaks per cell across 210s for each indicated viral concentration of Kir2.1 and NavD (*n* = 150 cells per condition). Differences were analyzed using Dunnett’s multiple comparison test on the Ca^2+^ transient frequency (S5d Fig) compared to the control (Kir2.1 0):(NavD 0) VG/*μ*l. The experimental trials that were statistically greater were marked (\*\*\*\**p* ≤ .0001) and (\*\*\**p* ≤ .0002).

The total cumulative peaks (S5A Fig) were greatest in the condition with a viral titer of (Kir2.1 1e8):(NavD 1e8) VG/*μ*L. To understand which viral titer impacted the cells we examined the cumulative number of Ca^2+^ transients divided by the total number of cells (Fig 7B). Using the data from the total number of peaks per ROI (S5d Fig) we ran Dunnett’s test (S1 Table) to examine if there was a statistical significance between the conditions. The experimental trials that were statistically greater than the control were marked (\*\*\*\**p* ≤ .0001, \*\*\**p* ≤ .0002) on the graph (Fig 7B).

It appears that using comparable amounts of the two channels (Kir2.1 1e8) and (NavD le8) VG/uL evoked a significant response because there is a 40% chance that out of 150 cells a significant peak will occur in any given cell in 210s. Of the cells that were suprathreshold (S5d Fig) there is a 47% chance of a second peak within ~50s (S5F Fig) after the initial peak. The mean event width decreased significantly in Kuhl-CKNa cells with (1e8 Kir2.1):(1e8 NavD) VG/*μ*L and Cx-43 (S3 fig).

### Optimizing Activity with Connexin-43

To further improve activity connexin-43 was optimized by varying the viral titer. The following concentrations were held constant for Kir2.1, NavD, bPAC, CNG, R-GECO (S2 Table). The Kuhl-CKNaCx cells were transduced and ~48 hours later the cells were stimulated with 20ms of blue light at 0s, 130s, and 240s. Images were collected with 561nm excitation.

The total cumulative peaks (S6A Fig) was greatest in the condition with a viral titer of (CNG le5) VG/*μ*L. To understand which viral titer impacted the chance of a cell firing we examined the cumulative number of Ca^2+^ transients divided by the total number of cells (Fig 8). Using the data from the total number of peaks per ROI (S6D Fig) we ran Dunnett’s test (S1 Table) to examine if there was a statistical significance between the conditions compared to the control (Cx-43 0) VG/*μ*l. The experimental trials (Cx 1e5) (\*\*\**p* ≤ .0003) and (Cx lei) (\*\**p* ≤ .0062) had a 100% chance that out of 150 cells they would fire at least twice in 400s. Further, there is a 36% chance that the cells that were suprathreshold (S6D Fig) are likely to produce another Ca^2+^ peak ~112s after the initial fluorescence peak (S6F Fig). While (Cx 1e1) was significant we chose to use (Cx 1e5) because it is 20% more likely to produce a significant fluorescence peak.

**Fig 8.**
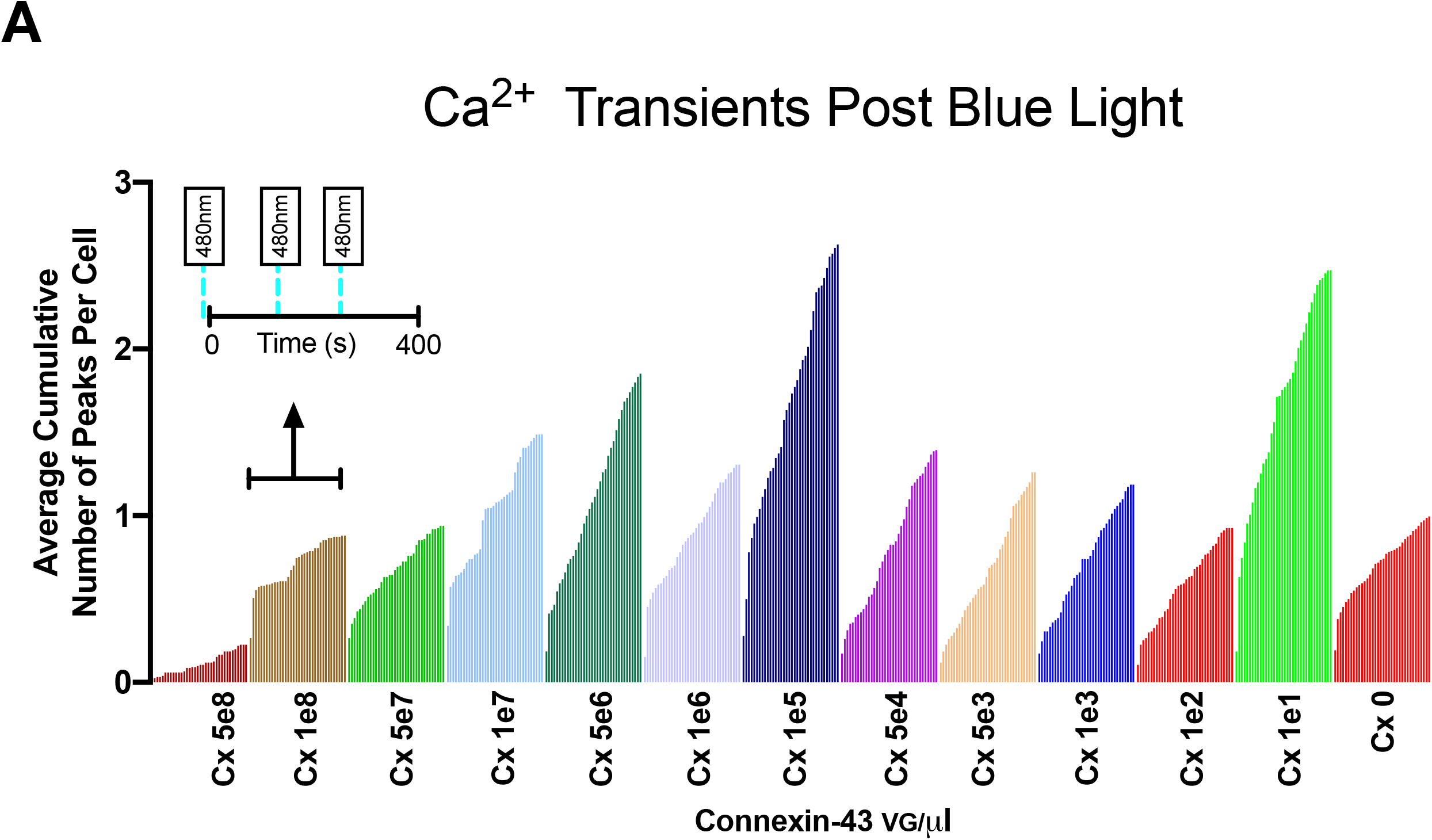
The gap junctions between cells using connexin-43 was optimized by varying levels of viral titer. The representative cartoon (Fig 7A) depicts all the components transduced into the experimental Kuhl-CKNaCx system (S2 Table). A) Shown are the average cumulative fluorescence peaks per cell across 400s for each indicated viral concentration of Cx-43 (*n* = 150 cells per condition).

### Optimizing Activity with CNG

The addition of NavD and connexin-43 had such a profound effect on the activity that we decided to re-optimize the CNG and HCN2 channel. To optimize activity with CNG the viral titer was varied and the following concentrations were held constant for Kir2.1, NavD, bPAC, Cx-43, and R-GECO1 (S2 Table). The Kuhl-CKNaCx cells were transduced and the following day the cells were stimulated with 20ms of blue light at 130s and 240s. Images were collected with 561nm excitation before and following each blue light stimulus.

Upon activation of bPAC there were various responses in each condition that are fully quantified in (S7 Fig). The total cumulative peaks (S7A Fig) was greatest in the condition with a viral titer of (CNG 1e5) VG/*μ*L. To understand which viral titer impacted the chance of a cell firing we examined the average cumulative number of Ca^2+^ transients divided by the total number of cells (Fig 9A). Using the data from the total number of peaks per ROI (S7A Fig) we used Dunnett’s test (S1 Table) to examine if there was a statistical significance between the conditions. The experimental trials (CNG 1e2 to 1e5) VG/*μ*L (\*\*\*\**p* ≤ .0001) were significantly greater than the control (CNG 0) VG/*μ*l and had a 100% chance that out of 150 cells they would fire at least once in 360s. Further, in Kuhl-CKNaCx (CNG le3) there is a 40% chance that of the cells that fired (S7D Fig) are likely to fire again ~80s after the initial fluorescence peak (S7F Fig).

**Fig 9.**
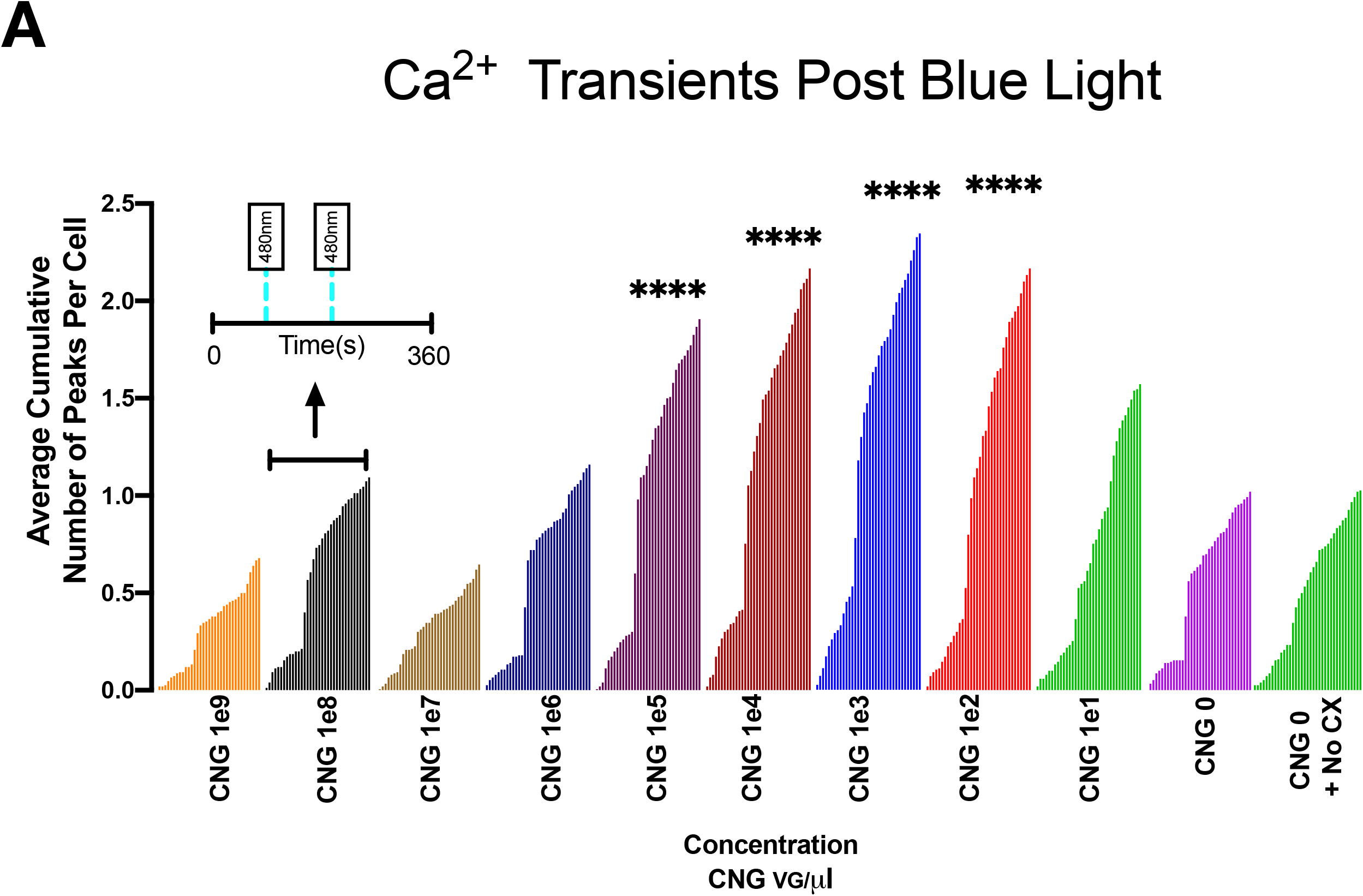
The amount of CNG channel was optimized. The representative cartoon (Fig 7A) depicts all the components transduced into the experimental Kuhl-KCNaCx system (S2 Table). A) Shown are the average cumulative fluorescence peaks per cell for each indicated viral concentration of CNG (*n* = 150 cells per condition). Differences were analyzed using Dunnett’s multiple comparison test on the Ca^2+^ transient frequency (S7D Fig) compared to the control (CNG 0) VG/*μ*l. The experimental trials that were statistically greater than the control were marked (\*\*\*\**p* ≤ .0001) on the graph.

### Optimizing Activity with HCN2

The HCN2 ion channel can be exchanged for the modified CNG channel. To optimize activity the viral titer for HCN2 was varied and the following concentrations held constant for Kir2.1, NavD, bPAC, Cx-43 and R-GECO1 (S2 Table). The Kuhl-HKNaCx cells were transduced and ~48 hours following the cells were stimulated with 20ms of blue light at 120s and 200s. Images were collected before and following each blue light stimulus with 561nm excitation.

Upon activation of bPAC there were somewhat varied responses in each condition that are fully quantified in (S8 Fig). The total cumulative peaks (S8A Fig) was greatest in the condition with a viral titer of (HCN2 1e2) VG/*μ*L. To understand which viral titer impacted the chance of a cell firing we examined the average cumulative number of Ca^2+^ transients divided by the total number of cells (Fig 10B).

**Fig 10.**
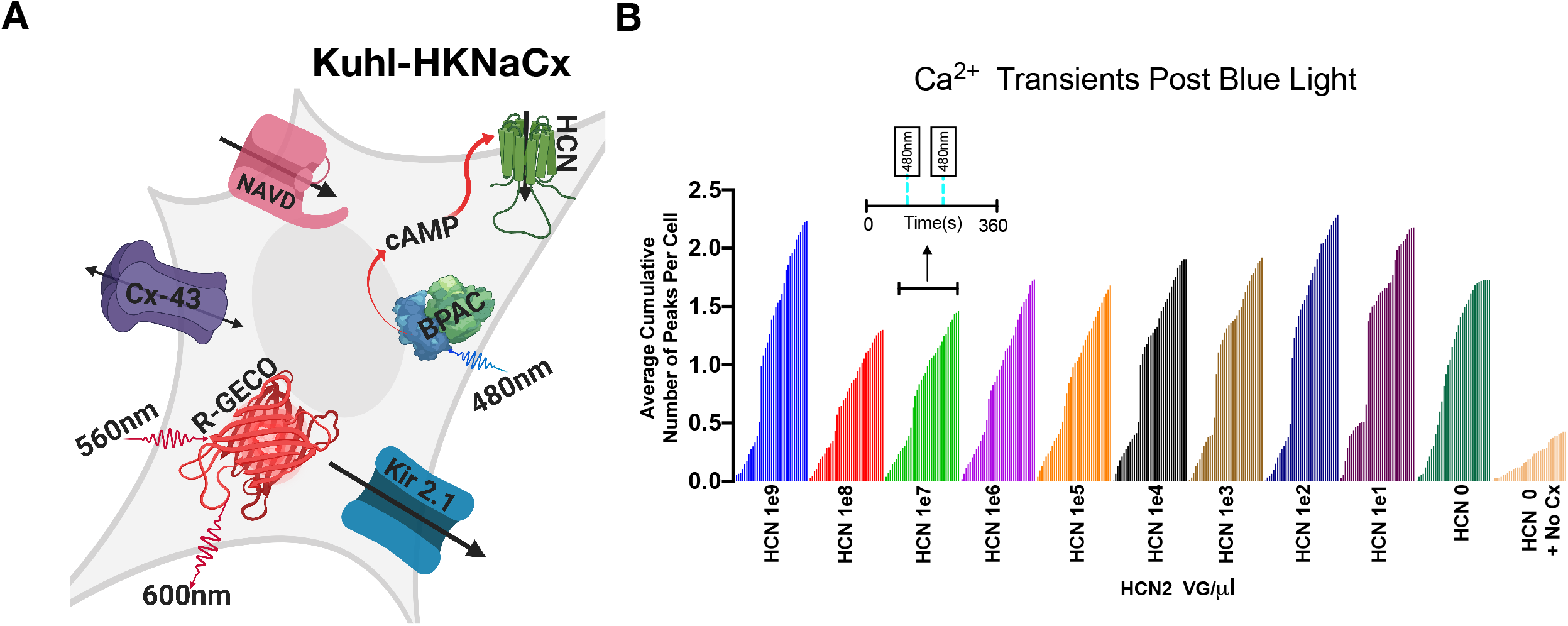
HCN2 can be exchanged for the modified CNG channel. A) Cartoon depicting all the components transduced into the experimental Kuhl-HKNaCx cell (S2 Table). B) Shown are the average cumulative fluorescence peaks per cell across 360s for each indicated viral concentration of HCN2 (*n* = 150 cells per condition).

Using the data from the total number of peaks per ROI (S8 Fig) we used Dunnett’s test (S1 Table) to examine if there was a statistical significance between the conditions. The experimental trials (HCN2 1e1, 1e2, and 1e9) VG/*μ*L (\**p* ≤ .0001) were significantly greater than the control (HCN2 0) VG/*μ*l. Further in (HCN2 le2) Kuhl-HKNaCx cells there is a 41% chance that out of 150 cells a significant peak will occur and of the cells that were above threshold (S8D Fig) are likely to fire ~77s after the initial fluorescence peak (S8F Fig).

It appears that the optimal concentration is (HCN2 1e2) VG/*μ*L but there appeared to be a range between (HCN2 1e1 to 1e2) VG/*μ*L (Fig 10B). The controls still display some random activity even without the HCN2 channel due to having the Kir2.1, NavD, and Cx-43 channel expressed. However, unlike the CNG control data (Fig 9A) the activity seems to be dependent on the Cx-43 being present as well as the Kir2.1 and NavD channel.

The behavior of HCN2 was visually different from CNG. While both wells have random activity, there is a brief somewhat coordinated activity after a blue light stimulus (S7C and S8C Figs). S2 and S3 Movies show the movement of CNG and HCN2. CNG has a Ca^2+^ bursting rosette pattern following a stimulus, while HCN2 seems to have a spreading wave-like pattern.

### A KUHL-KCNaCx

The Kuhl-KCNaCx cells show remarkable, consistent activity when stimulated with blue light. One can image using them to screen prototype activity sensors, new GECI and GEVI biosensors. Such a screen, however, would depend critically on consistent well to well activity. To measure how consistent the activity is we measured the activity in 14 wells of cells transduced with an optimized amount of each channel. The representative cartoon (Fig 11A) depicts all the components transduced into the Kuhl-KCNaCx cell. The Kuhl-KCNaCx cells (S2 Table) were transduced and ~48 hours later the cells were imaged for 25 minutes and stimulated with 20ms of blue light at 500s and 1000s. Images were collected before and after each blue light stimulus with 561nm excitation for 4 wells over 25 minutes (Fig 11A) or 10 wells for 15 minutes (S4 Movie).

**Fig 11.**
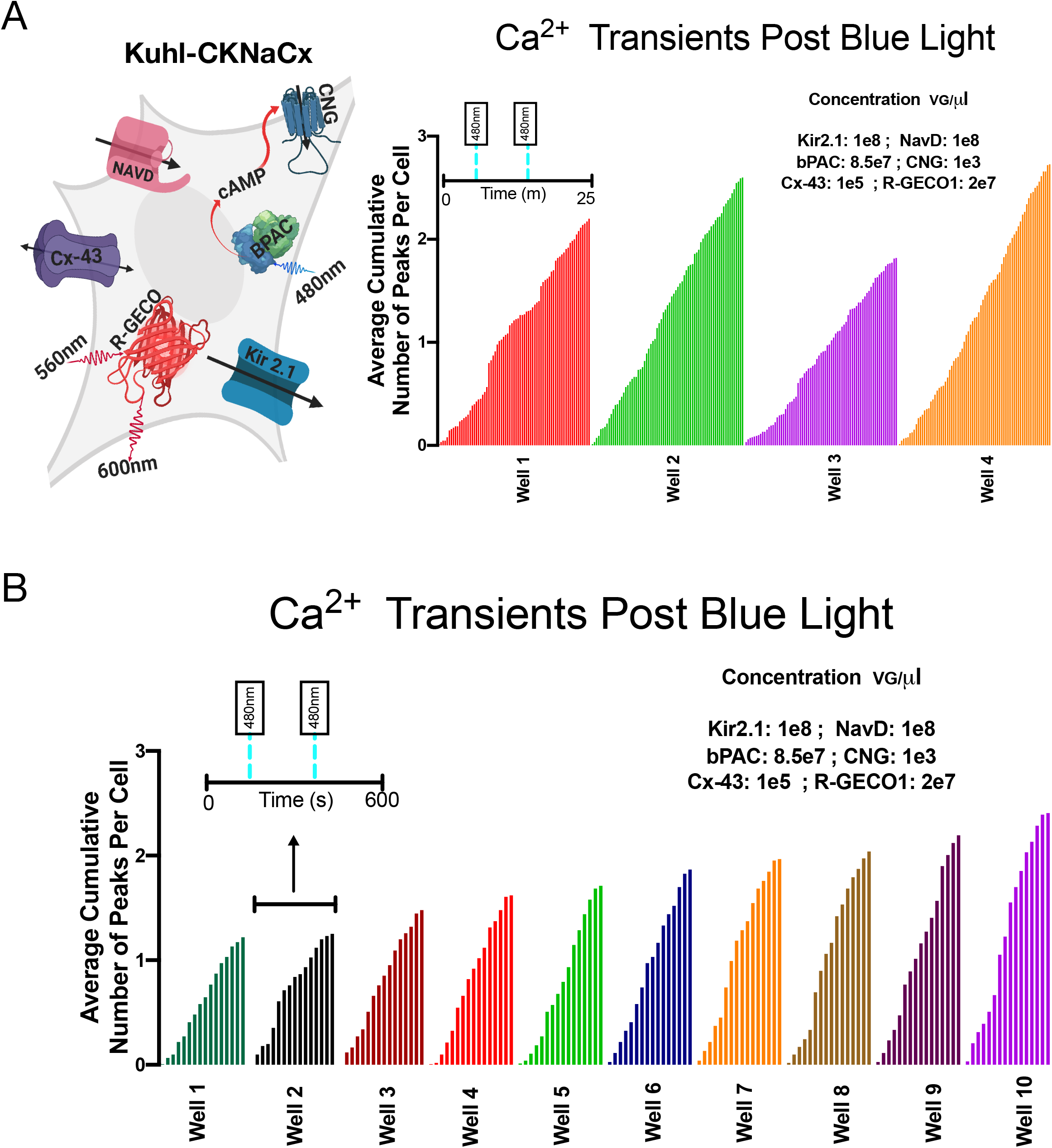
A Kuhl-CKNaCx - Consistent activity with stable viral concentration. Well to well average cumulative peak activity is fairly consistent. A) The representative cartoon depicts all the components transduced into the Kuhl-CKNaCx cell (S2 Table). The average cumulative number of peaks per cell over four trials in 25m. B) The average cumulative number of peaks per cell over 10 trials in 15m. Tukey’s multiple comparison test was run on the Kuhl-CKNaCx trials and found no significant difference between the wells.

The well to well average cumulative peak activity (Fig 11A) has a mean of 2.4 ±.4. Taking the mean number of total peaks per trial by the mean total number of peaks that in Kuhl-CKNaCx cells 100% of the 600 cells will produce a significant peak and of the cells that were above threshold (S9F Fig) 50.8% are likely to produce a significant peak ~94s ±32 following.

Another 10 wells (Fig 11B) were imaged for 15 minutes and stimulated with 20ms of blue light at 198s and 400s. There appears to be increased coordinated activity between the cells following blue light stimulus (S10C Fig). The cells have an oscillating firing pattern (S10C Fig) that occurs and is consistent with the mean inter-event interval (S10F Fig) for each well. The well to well activity when measuring the average cumulative number of peaks per cell is fairly consistent with a mean of 1.8±.4. There is a 100% chance that out of 1,500 cells a significant peak will occur 1,500 times when imaged for 15 minutes and of the cells that were above threshold (S10F Fig) 29% are likely to produce a significant peak ~148s±89 following the initial peak. Tukey’s multiple comparison test found that there wasn’t a significant difference between the average cumulative number of peaks per cell for the 14 trials with Kuhl-CKNaCx.

### Comparing Green and Red Sensors in the Same Cell

While slow and laborious, an advantage of patch clamp fluorometry is that whole-cell voltage changes can be directly compared with fluorescence changes. The all-optical Kuhl cell approach does not have the advantage of such an independent measurement. However, we reasoned that different colored sensors could be simultaneously measured to arrive at a comparative measure. For example, a green sensor of known properties could be compared to a new one with red fluorescence. To test the feasibility of this approach, we used an optical splitter to compare the responses of R-GECO1 and G-GECO1 within the same cell.

The Kuhl-KCNaCx cells were transduced and 48 hours following the cells were imaged. In figure 12 an image created by the OptoSplit II is shown where the green and red emission from both sensors are split and imaged on two different portions of the EMCCD camera(S5 Movie). The fluorescence response from both sensors is plotted in figure 12 (Fig 12B). Previous photophysical characterization of G-GECO1 and R-GECO1 show that there are relatively large differences between the two sensors. G-GEC01 has a faster K_on_ and R-GECO1 has a faster K_off [8]_. G-GECO1 has a greater K_d, kinetic_ with a difference of 347(nm)^1^. G-GECO1 also is ~150x brighter than R-GECO1 [8]. This simple analysis shows that differences can be easily measured in Figure 12 (Fig 12B) where the Tau_on/off_ of both signals was measured. R-GECO1 Tau_on_ was consistently slower than G-GECO1 and R-GECO1 Tau_off_ was consistently faster than G-GECO1 Tau_off_.

**Fig 12.**
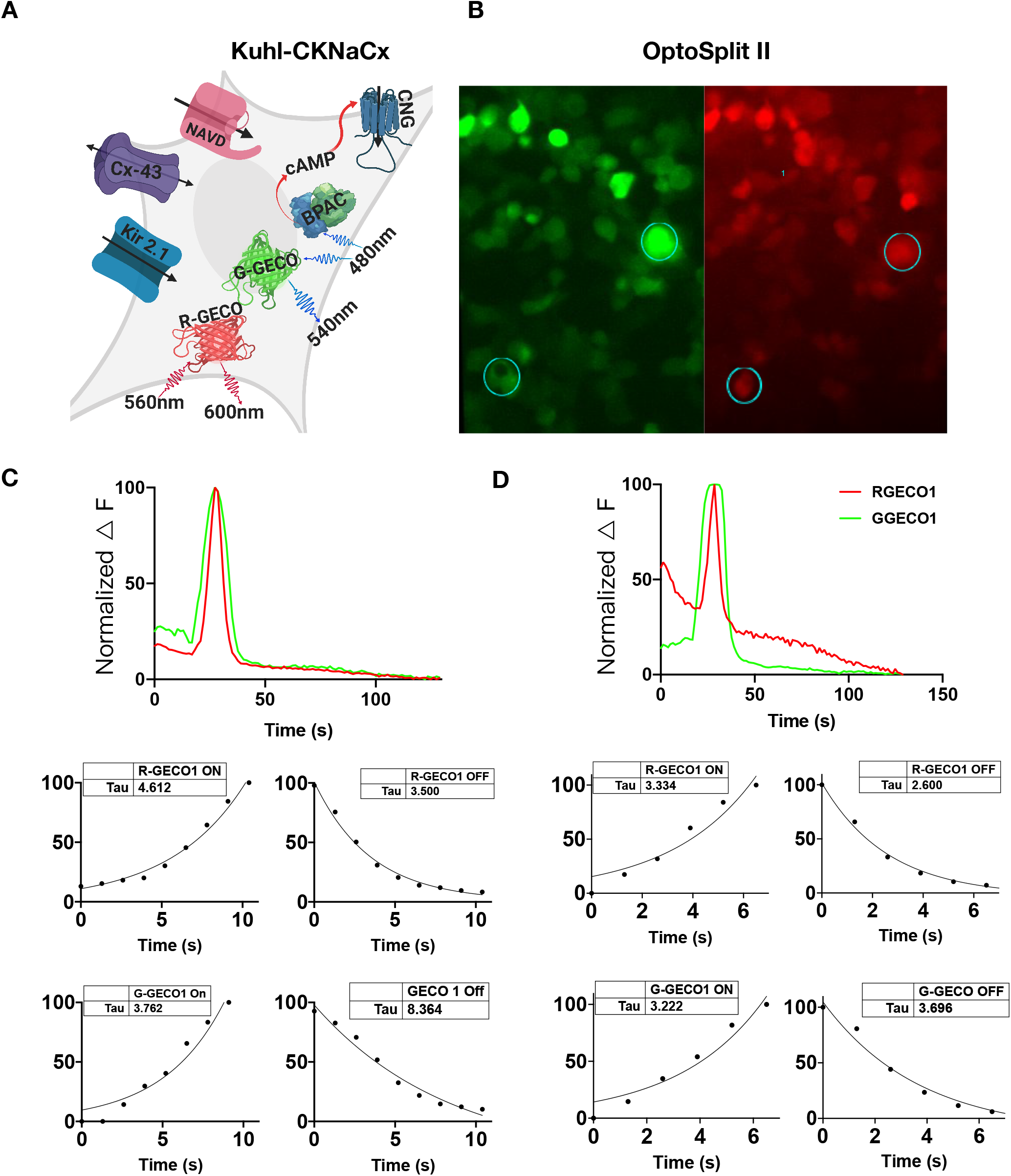
Both red and green Ca^2+^ sensors can be recorded from the same cell and imaged simultaneously using an OptoSplit II. A) Cartoon depicting all the components transduced into the Kuhl-CKNaCx cell. bPAC was transfected at a low level so that the cells could be imaged with green light continuously. B) The cells were simultaneously imaged with green and red wavelengths for 150s. The image is of the same cells in two different wavelengths and ROIs from both the G-GECO1 and R-GECO1 were obtained (circles and numbers). Representative traces from the ROIs show that both G-GECO1 and R-GECO1 fluorescence are visible and their respective responses can be compared and plotted over time. The difference between G-GECO1 and R-GECO1 can be seen because it has a greater intensity change (150x) in fluorescence to Ca^2+^ concentrations and has a ~2 fold increase in the speed of its kinetics (K_d_).

### Green Voltage Sensors Can be Screened in Kuhl-CKNaCx Cells

ArcLight is a downward going voltage sensor, meaning that it gets brighter upon the hyperpolarization of the membrane and dimmer at depolarization. It is pH sensitive and has been demonstrated to work in single-cell fly neurons but requires high-intensity illumination to achieve an adequate signal-to-noise ratio [39].

The following concentrations were held constant for Kir2.1, NavD, bPAC, Cx-43, CNG, ArcLight (S2 Table). The Kuhl-CKNaCx cells were transduced and ~48 hours following the cells were imaged with continuous 480nm excitation. Slow depolarizations can be seen in the Kuhl-CKNaCx cell system (Fig 13) with a reasonable SNR of 5.5 ±4 averaged from 15 representative traces. We believe that the Kuhl-CKNaCx system shows promise for screening voltage sensor prototypes(S6-S8 Movie).

**Fig 13.**
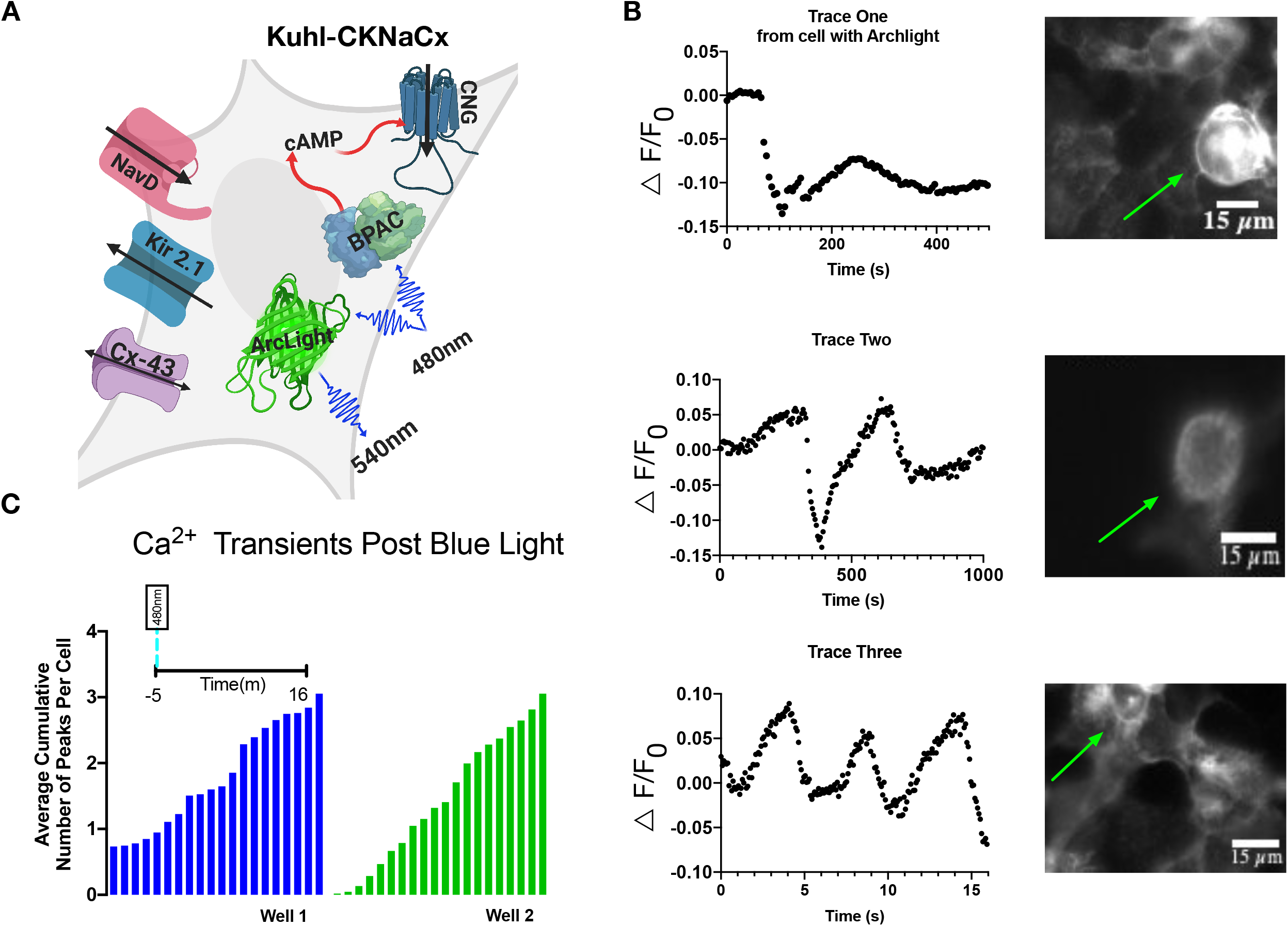
The Kuhl-CKNaCx system undergoes voltage changes that are visible with ArcLight. A) Cartoon depicting all the components transduced into the Kuhl-CKNaCx cell. bPAC was stimulated continuously while exciting Arclight. B) Blue light activation of bPAC caused a slow depolarization that occurred over a few seconds. Representative traces from the ROIs show that voltage changes are visible using ArcLight. Images of ArcLight fluorescence response over time.The green arrows annotate which cells were used for the fluorescence traces. C) Two separate wells that were imaged are shown.

## Discussion

This Kuhl cell system was designed to be customizable and there are several important considerations to take into account when customizing for a particular activity sensor. Multiple channels could be exchanged for the ones used in this study. For example, a faster voltage-gated sodium channel would most likely create a faster action potential in the cells.

The Kuhl cells can potentially be used in high throughput screens for better activity biosensors. The throughput of any screen, however, depends on the time budget. Here we arbitrarily picked fluorescence peaks that exceeded the SNR of 26. That is, the peak was 26 times larger than the standard deviation of the baseline noise. Using this threshold, we found that out of 150 cells, imaging for 8Os±3O was sufficient to detect 100 cells that would produce a Ca^2+^ peak above threshold. Recording the activity in a well for 80s would eliminate false negatives. The imaging time, plus the plate translation time, should result in a time budget of roughly 100 minutes per 96 well plate. While 400 to 500 wells a day is low throughput, it is considerably better than the manual patch clamp fluorometry used today.

We have shown that screening both red and green GECIs simultaneously in the Kuhl-CKNaCx system is possible, and that one sensor can be used to benchmark the other. Individual cells’ dual GECI response can be measured for its brightness, kinetics (Tau), SNR, and sensitivity. In theory, if a lower objective lens was used, an average of the field of view for the mutant sensor’s fluorescence response could be easily compared to the average response of the benchmark sensor in that same well.

The improvement of the genetically encoded voltage indicators - GEVIs - has been slow due to limitations in the throughput involved in manual, whole-cell patch clamp fluorometry [12,40]. The system described here will enable investigators to screen without field stimulation or any manual intervention. Voltage changes along the membrane of the HEK293 cells are visible and can be tracked in the Kuhl-CKNaCx cells using the GEVI ArcLight, since the action potential in these cells is quite slow, lasting seconds ~4s. We believe that because we were able to screen and benchmark Ca^2+^ indicators simultaneously that this should also work for voltage sensors. The prototype sensors could be expressed in Kuhl cells that also express a benchmark sensor such as GCAMP7f or Voltron [2,41].

## Methods

### Growth conditions for HEK293 cells

A line of HEK293 were cultured in DMEM, 10% FBS, and 1% Penicillin (100 U/mL). For experiments, HE293 cells were plated in 96 well plastic plates at 100ml per well (~3-5 x10^4^ cells/mL).

### Plasmids

The HCN2 plasmid was kindly donated by Joan Lemire at Tufts University from Dr. Michael Levin’s lab. The NavRosDg217a was reverse translated using a human codon preference and synthesized as a gBblock by IDT (Coralville, Iowa). The cyclic nucleotide-gated olfactory channel (from *Rattus norvegicus*) has the following mutations, *δ*61-90/C460W/E583M and was synthesized as a gBblock by IDT. [42]. pGEMTEZ-Kir2.1 was a gift from Richard Axel, Joseph Gogos & Ron Yu (Addgene plasmid #32641).

### Baculovirus

The channels Kir2.1, CNG, HCN2 and NavRosDg217a were packaged in BacMam (Montana Molecular, Bozeman, MT). The virus titers were: Kir2.1 3.6×10^10^ VG/mL, bPAC 8.59 x 10^10^ VG/mL, CNG 3.55×10^10^ VG/mL, and NavRosDg217a 7.26× 10^10^ VG/mL. The following bio-sensors in baculovirus were obtained from Montana Molecular: R-cADDis (titer 5.06×10^10 VG/mL); R-GECO (2×10^10^ VG/mL); G-GECO (9.17×10^10^VG/mL); and Arclight (2×10^10^VG/mL). Supplemental figure 12 depicts each experimental condition and the final viral concentrations used.

### Transduction

HEK-293 cells were prepared and transduced following the manufacturer’s recommended protocol. Briefly, HEK293 cells were plated in 96 well plastic plates (48 x10^3^ cells/well) and transduced with the appropriate mix of viruses and the HDAC inhibitor sodium butyrate (2mM final concentration). Two days later, the media was exchanged with PBS, tyrodes buffer, or Fluorobrite. The best results were consistently obtained with PBS.

### An ASI modular microscope

The experiments (Figs 1–11 and 13) were conducted using wide field imaging on an ASI-XYZ stage fitted with a modular infinity microscope. The objective imaged onto the detector chip of a Hamamatsu ORCA-Flash 4.0 scientific-CMOS camera. The images were collected using either MATLAB scripts that controlled the camera, ASI stage, SH1-Thorlabs shutter, and ThorLabs DC4100 Four Channel-LED Driver, or by using μManager [43]. A custom illumination system was arranged. Briefly, a dichroic mirror was positioned at the entrance of the microscope to combine blue light LED illumination with 561 laser illumination. An SH1-Thorlabs shutter was used to create brief illumination from a blue ThorLABS LED (488nm) for rapid stimulation of bPAC. At 100% the blue light illumination was 70mW/cm^2^. The yellow illumination was provided with a Sapphire laser (561nm, 50mW, Coherent). The laser beam was steered with two mirrors (arranged in a periscope) into an entrance aperture of the ASI microscope. Before entering the microscope, the laser beam passed through a 50° diffuser (ED1 C50 MD, Thorlabs) placed in the focus of an f = 20 mm aspheric lens (ACL2520U, Thorlabs) that collimated the beam for further traveling into the microscope.

### Optosplitter

To simultaneously collect images of both G-GECO and R-GECO, we used an optical splitter (CAIRN optosplit II, Faversham Kent ME13 8UP). This makes it possible to simultaneously collect both the green and red images on a single camera. An Olympus 1X81 was fitted with LED illumination (470 nM, 565 nM, ThorLabs) where the power to the LEDs can be independently adjusted. A Pinkel filter set was used to simultaneously image both G-GECO and R-GECO (GFP/DsRed-2X-A-000, Semrock, Rochester, New York). The CAIRN splitter was positioned in between the microscope and EMCCD camera such that a dichroic filter (561 nM, Semrock, Rochester, New York) split the green and red emission into two optical paths. The red emission passed through an additional band pass filter (617/70, Semrock, Rochester, New York) to eliminate any potential green emission from the G-GECO.

### Analysis

Image data was stored in a Z-stack tiff file and loaded into the FIJI distribution of the ImageJ software [44]. The initial frame of the stack was eliminated from all analysis to correct for photoactivation of R-GECO1 upon bluelight stimulation of bPAC [45]. The cells were selected using a freehand ROI surrounding the cell of interest. The average total intensity pixel value within the ROI for each frame was collected using the time series analyzer plugin and saved as a .csv file. The raw fluorescence trace data was loaded into MATLAB.

The analysis was done in MATLAB using the Find Peaks function in the Signal Processing Toolbox. The raw fluorescence traces were rescaled from zero to 10,000. The data was smoothed using ‘sgolay’ function. The main peak prominence threshold was set to 50.5 with an SNR of 26±10. The Signal-to-noise ratio was defined as the average of the peaks amplitudes that were defined using the peak finder thresholder by the standard deviation (S.D.) of the signal before blue light stimulus. We examined the SNR for Kuhl-CKNaCx cells over 5 individual trials which were averaged over 15 trials. The FindPeaks function returns a vector with the local maxima (peaks) from the ROI trace data. The Peak Finder in MATLAB marks each fluorescence peak by its location in time and records the number of peaks per ROI. In addition, it returns the widths of the peak and the prominence of the peak. The time that a fluorescence peak occurred during imaging was recorded. The inter-event interval is the absolute difference between each consecutive peak. The FWHM and inter-event interval time was adjusted for exposure times. The total cumulative number of peaks for an entire trial was measured. Each fluorescence peak was measured from zero to the maximum peak intensity (normalized F) using the MATLAB Peak Finder. The time and total quantity of a fluorescence peak in a ROI were recorded. To determine the chance that a cell will fire twice was determined by (total number of peaks for a well/total number of points collected from Inter-Peak interval)

## Acknowledgements

The authors thank the following people: Paul Tewson for HEK293 cells. Elsa Roush for multiple 96 well plates of transduced HEK293 cells. Scott Martinka for excellent high titer virus. Anne Marie. Rosana Molina. Steven Hoffman, Dr. Jamie Mazer, Dr. Kaspar Podgorski, and Nadav Oakes for MATLAB assistance. Joan Lemire at Tufts University for HCN2 plasmid. Mikhail Drobizhev was a constant source of advice and optical expertise. The reviewers for their invaluable suggestions.

## Supporting Information

**S1 Fig.** Fluorescence trace of cells with Kir2.1, CNG, R-GECO1 (Kuhl-CK). Without bPAC blue light stimulus is ineffective at creating a response within cells.

**S2 Fig. Representative traces.** Raw fluorescence traces from Kuhl-CK AND Kuhl-HK. 20s of blue light stimulus leads to varied fluctuating responses in each cell.

**S3 Fig. Introduction of NavD to Kuhl and Cx-43 increases the speed of the Ca^2+^ transients.** A) The most significant wavelengths from each optimized set was included in the data to show the increase in Ca^2+^ transient speed.

**S4 Fig. Features of the [Kir2.1]:[NavD] optimized Kuhl-CKNa cells and their Ca^2+^ transients.** A) The total peaks for each Ca^2+^ transient within a trial was recorded, and the experiment with [1e8 Kir2.1]:[5e8 NavD] VG/*μ*L had the greatest recorded peaks. B) The □F is the difference between the baseline fluorescence and the maximum fluorescence of each peak. The mean and S.E.M of the Ca^2+^ transients in [Kir2.1]:[NavD] optimized Kuhl-CKNa cells C) Blue line indicates point of 20s blue light stimulus. The time of each fluorescence peak was recorded and is indicated by a black dot. D) The total number of Ca^2+^transients per cell (ROI) for experimental [Kir2.1]:[NavD] optimized Kuhl-CKNa cells. E) Representative duration of elevated R-GECO1 fluorescence over time per Ca^2+^ transient. Black dot indicates the Ca^2+^transients FWHM for each peak. F) Black dot indicates the time between Ca^2+^ transient events per cell. Red bars indicate mean and ± S.E.M (*n* = 150 cells per condition).

**S5 Fig. Features of the [Kir2.1]:[NavD] optimized Kuhl-CKNaCx cells and their Ca^2+^ transients.** A) The total peaks for each Ca^2+^ transient within a trial was recorded, and the experiment with [1e8 Kir2.1]:[1e8 NavD] VG/*μ*L had the greatest recorded peaks. B) The □F is the difference between the baseline fluorescence and the maximum fluorescence of each peak. The mean and S.E.M of the Ca^2+^ transients in [Kir2.1]:[NavD] optimized Kuhl-CKNaCx cells C) Blue line indicates point of 20s blue light stimulus. The time of each fluorescence peak was recorded and is indicated by a black dot. D) The total number of Ca^2+^transients per cell (ROI) for experimental [Kir2.1]:[NavD] optimized Kuhl-CKNaCx. E) Representative duration of elevated R-GECO1 fluorescence overtime per Ca^2+^transient. Black dot indicates the Ca^2+^transients FWHM for each peak. F) Black dot indicates the time between Ca^2+^ transient events per cell. Red bars indicate mean and ± S.E.M (*n* = 150 cells per condition).

**S6 Fig. Features of the Cx-43 optimized Kuhl-CKNaCx cells and their Ca^2+^ transients. Corresponding figure 8.** A) The total peaks for each Ca^2+^ transient within a trial was recorded, and the experiment with [Cx 1e5] VG/*μ*L had the greatest recorded peaks. B) The □F is the difference between the baseline fluorescence and the maximum fluorescence of each peak. The mean and S.E.M of the Ca^2+^ transients in Cx-43 optimized Kuhl-CKNaCx cells. C) Blue line indicates point of 20s blue light stimulus. The time of each fluorescence peak was recorded and is indicated by a black dot. D) The number of Ca^2+^ transients per cell (ROI) for Cx-43 optimized Kuhl-CKNaCx cells. E) Representative duration of elevated R-GECO1 fluorescence over time per Ca^2+^transient. Black dot indicates the Ca^2+^transients FWHM for each peak. F) Black dot indicates the time between Ca^2+^ transient events per cell. Red bars indicate mean and ± S.E.M (*n* = 150 cells per condition).

**S7 Fig: Features of the CNG optimized Kuhl-CKNaCx cells and their Ca^2+^ transients. Corresponding figure 9.** A) The total peaks for each Ca^2+^ transient within a trial was recorded, and the experiment with [CNG 1e3] VG/*μ*L had the greatest recorded peaks. B) The □F is the difference between the baseline fluorescence and the maximum fluorescence of each peak. The mean and S.E.M of the Ca^2+^ transients in CNG optimized Kuhl-CKNaCx cells C) Blue line indicates point of 20s blue light stimulus. The time of each fluorescence peak was recorded and is indicated by a black dot. The wells without CNG and Cx were very interesting because it appears that without blue light there is a D) The total number of Ca^2+^ transients per cell (ROI) for experimental CNG optimized Kuhl-CKNaCx cells. E) Representative duration of elevated R-GECO1 fluorescence over time per Ca^2+^transient. Black dot indicates the Ca^2+^transients FWHM for each peak. F) Black dot indicates the time between Ca^2+^ transient events per cell. Red bars indicate mean and ± S.E.M (*n* = 150 cells per condition).

**S8 Fig. Features of the HCN2 optimized Kuhl-HKNaCx cells and their Ca^2+^ transients. Corresponding figure 10.** A) The total peaks for each Ca^2+^ transient within a trial (T=300s) was recorded. The experiment with [HCN2 1e2] VG/*μ*L had the greatest recorded peaks. B) The ØF is the difference between the baseline fluorescence and the maximum fluorescence of each peak. The mean and S.E.M of the Ca^2+^ transients in HCN2 optimized Kuhl-HKNaCx cells C) Blue line indicates point of 20s blue light stimulus. The time of each fluorescence peak was recorded and is indicated by a black dot. D) The total number of Ca^2+^transients per cell (ROI) for experimental HCN2 optimized Kuhl-HKNaCx. E) Representative duration of elevated R-GECO1 fluorescence over time per Ca^2+^transient. Black dot indicates the Ca^2+^transients FWHM for each peak. F) Black dot indicates the time between Ca^2+^ transient events per cell. Red bars indicate mean and ± S.E.M (*n* = 150 cells per condition).

**S9 Fig. Features of the optimized Kuhl-CKNaCx cells and their Ca^2+^ transients. Corresponding figure 11a.** A) The total peaks for each Ca^2+^ transient within a trial (T=25m) was recorded. B) The □F is the difference between the baseline fluorescence and the maximum fluorescence of each peak. The mean and S.E.M of the Ca^2+^ transients in optimized Kuhl-CKNaCx cells C) Blue line indicates point of 20s blue light stimulus. The time of each fluorescence peak was recorded and is indicated by a black dot. D) The total number of Ca^2+^transients per cell (ROI) for optimized Kuhl-CKNaCx cells. Representative duration of elevated R-GECO1 fluorescence over time per Ca^2+^transient. Black dot indicates the Ca^2+^transients FWHM for each peak. F) Black dot indicates the time between Ca^2+^ transient events per cell. Red bars indicate mean and ± S.E.M (*n* = 150 cells per condition).

**S10 Fig. Features of the optimized Kuhl-CKNaCx cells and their Ca^2+^ transients. Corresponding figure 11b.** A) The total peaks for each Ca^2+^ transient within a trial (T=25m) was recorded. B) The □F is the difference between the baseline fluorescence and the maximum fluorescence of each peak. The mean and S.E.M of the Ca^2+^ transients in optimized Kuhl-CKNaCx cells C) Blue line indicates point of 20s blue light stimulus. The time of each fluorescence peak was recorded and is indicated by a black dot. D) The total number of Ca^2+^ transients per cell (ROI) for optimized Kuhl-CKNaCx. Representative duration of elevated R-GECO1 fluorescence over time per Ca^2+^ transient. Black dot indicates the Ca^2+^ transients FWHM for each peak. F) Black dot indicates the time between Ca^2+^ transient events per cell. Red bars indicate mean and ± S.E.M (*n* = 150 cells per condition).

**S1 Table. Dunnett’s Multiple comparisons test.** The table shows all test run using Dunnetts where stated in the manuscript.

**S2 Table. Experimental setup for each figure.** The table shows the experimental set up, the final viral concentration used, and their corresponding figures.

**S1 Movie. Bright Subcellular Flashes of Ca2+.** The authors do not know what these come from.

**S2 Movie. The temporal/spatial movement of CNG in Kuhl-CKNaCx cells.** Rose like patterns.

**S3 Movie. The temporal/spatial movement of HCN2 in Kuhl-HKNaCx cells.** Wave-like spreading pattern.

**S4 Movie. A well from the Kuhl-CKNaCx 10 well experiment.** Cells imaged for 15 minutes with robust, repeatable activity.

**S5 Movie. G-GECO1 and R-GECO1 can be imaged simultaneously.** An image created by the OptoSplit II is shown where the green and red emission from both sensors are split and imaged on two different portions of the EMCCD camera

**S6 Movie. Whole well image of ArcLight activity.** Activity of Kuhl-CKNaCx cells with the GEVI ArcLight.

**S7 Movie. Combined images of ArcLight activity.** Activity of Kuhl-CKNaCx cells with the GEVI ArcLight from the same well.

**S8 Movie. Combined images of ArcLight activity.** Activity of Kuhl-CKNaCx cells with the GEVI ArcLight from the same well.

## References

1. Stierl M, Stumpf P, Udwari D, Gueta R, Hagedorn R, Losi A, et al. Light modulation of cellular cAMP by a small bacterial photoactivated adenylyl cyclase, bPAC, of the soil bacterium Beggiatoa. J Biol Chem. 2011;286:1181–1188.

2. Kerruth S, Coates C, Dürst CD, Oertner TG, Török K. The kinetic mechanisms of fast-decay red-fluorescent genetically encoded calcium indicators. J Biol Chem. 2019;294: 3934–3946.

3. Huang L, Knoblich U, Ledochowitsch P, Lecoq J, Clay Reid R, de Vries SEJ, et al. Relationship between spiking activity and simultaneously recorded fluorescence signals in transgenic mice expressing GCaMP6. bioRxiv. 2019. p. 788802. doi:10.1101/788802

4. Agetsuma M, Matsuda T, Nagai T. Methods for monitoring signaling molecules in cellular compartments. Cell Calcium. 2017;64:12–19.

5. Bruton J, Cheng AJ, Westerblad H. Measuring Ca2+ in Living Cells. Adv Exp Med Biol. 2020;1131: 7–26.

6. Kim D, Svoboda K, Looger L, Schreiter E. Genetically encoded calcium indicators and methods of use. US Patent. 20190153067:A1, 2019. Available: https://patentimages.storage.googleapis.com/48/48/53/5bb27ffa25d100/US20190153067A1.pdf

7. Podgorski K, Ranganathan G. Brain heating induced by near-infrared lasers during multiphoton microscopy. J Neurophysiol. 2016;116:1012–1023.

8. Zhao Y, Araki S, Wu J, Teramoto T, Chang Y-F, Nakano M, et al. An expanded palette of genetically encoded Ca^2+^ indicators. Science. 2011;333:1888–1891.

9. Qian Y, Piatkevich KD, Mc Larney B, Abdelfattah AS, Mehta S, Murdock MH, et al. A genetically encoded near-infrared fluorescent calcium ion indicator. Nat Methods. 2019;16:171–174.

10. Dana H, Sun Y, Mohar B, Hulse BK, Kerlin AM, Hasseman JP, et al. High-performance calcium sensors for imaging activity in neuronal populations and microcompartments. Nat Methods. 2019;16: 649–657.

11. Ibraheem A, Yap H, Ding Y, Campbell RE. A bacteria colony-based screen for optimal linker combinations in genetically encoded biosensors. BMC Biotechnol. 2011;ll: 105.

12. Park J, Werley CA, Venkatachalam V, Kralj JM, Dib-Hajj SD, Waxman SG, et al. Screening fluorescent voltage indicators with spontaneously spiking HEK cells. PLoS One. 2013;8: e85221.

13. Ding Y, Li J, Enterina JR, Shen Y, Zhang I, Tewson PH, et al. Ratiometric biosensors based on dimerization-dependent fluorescent protein exchange. Nat Methods. 2015;12: 195–198.

14. Alford SC, Abdelfattah AS, Ding Y, Campbell RE. A fluorogenic red fluorescent protein heterodimer. Chem Biol. 2012;19: 353–360.

15. Zhao Y, Abdelfattah AS, Zhao Y, Ruangkittisakul A, Ballanyi K, Campbell RE, et al. Microfluidic cell sorter-aided directed evolution of a protein-based calcium ion indicator with an inverted fluorescent response. Integr Biol. 2014;6: 714–725.

16. Shen Y, Dana H, Abdelfattah AS, Patel R, Shea J, Molina RS, et al. A genetically encoded Ca2+ indicator based on circularly permutated sea anemone red fluorescent protein eqFP578. BMC Biol. 2018;16: 9.

17. Hsu H, Huang E, Yang XC, Karschin A, Labarca C, Figl A, et al. Slow and incomplete inactivations of voltage-gated channels dominate encoding in synthetic neurons. Biophys J. 1993;65: 1196–1206.

18. Kirkton RD, Bursac N. Engineering biosynthetic excitable tissues from unexcitable cells for electrophysiological and cell therapy studies. Nat Commun. 2011;2: 300.

19. Zhang H, Reichert E, Cohen AE. Optical electrophysiology for probing function and pharmacology of voltage-gated ion channels. Elife. 2016;5. doi:10.7554/eLife.15202

20. Shah BS, Rush AM, Liu S, Tyrrell L, Black JA, Dib-Hajj SD, et al. Contactin associates with sodium channel Nav1.3 in native tissues and increases channel density at the cell surface. J Neurosci. 2004;24: 7387–7399.

21. Zhang H, Cohen AE. Optogenetic Approaches to Drug Discovery in Neuroscience and Beyond. Trends Biotechnol. 2017;35: 625–639.

22. Khan KH. Gene expression in Mammalian cells and its applications. Adv Pharm Bull. 2013;3: 257–263.

23. Rodin SN, Riggs AD. Epigenetic silencing may aid evolution by gene duplication. J Mol Evol. 2003;56: 718–729.

24. Kost TA, Condreay JP, Ames RS, Rees S, Romanos MA. Implementation of BacMam virus gene delivery technology in a drug discovery setting. Drug Discov Today. 2007;12: 396–403.

25. Bernal Sierra YA, Rost BR, Pofahl M, Fernandes AM, Kopton RA, Moser S, et al. Potassium channelbased optogenetic silencing. Nat Commun. 2018;9: 4611.

26. Tewson PH, Martinka S, Shaner NC, Hughes TE, Quinn AM. New DAG and cAMP Sensors Optimized for Live-Cell Assays in Automated Laboratories. J Biomol Screen. 2016;21: 298–305.

27. Hirano M, Takebe M, Ishido T, Ide T, Matsunaga S. The C-terminal region affects the activity of photoactivated adenylyl cyclase from Oscillatoria acuminata. Sci Rep. 2019;9: 20262.

28. Fesenko EE, Kolesnikov SS, Lyubarsky AL. Induction by cyclic GMP of cationic conductance in plasma membrane of retinal rod outer segment. Nature. 1985;313: 310–313.

29. Rich TC, Fagan KA, Nakata H, Schaack J, Cooper DMF, Karpen JW. Cyclic Nucleotide–Gated Channels Colocalize with Adenylyl Cyclase in Regions of Restricted Camp Diffusion. J Gen Physiol. 2000;116: 147–162.

30. Chen K, Zuo D, Wang S-Y, Chen H. Kir2 inward rectification-controlled precise and dynamic balances between Kir2 and HCN currents initiate pacemaking activity. FASEB J. 2018;32: 3047–3057.

31. Capel RA, Collins TP, Bose SJ, Rajasundaram S, Ayagama T, Zaccolo M, et al. cAMP signalling is required for the actions of IP3 on Ca2+-transients in cardiac atria and beating rate in sino-atrial node. bioRxiv. 2019. p. 694349. doi:10.1101/694349

32. Nguyen HX, Kirkton RD, Bursac N. Engineering prokaryotic channels for control of mammalian tissue excitability. Nat Commun. 2016;7: 13132.

33. Hille B. Ionic Channels of Excitable Membranes. Sinauer; 2001.

34. Yu CR, Power J, Barnea G, O’Donnell S, Brown HEV, Osborne J, et al. Spontaneous neural activity is required for the establishment and maintenance of the olfactory sensory map. Neuron. 2004;42: 553–566.

35. Woehler A, Wlodarczyk J, Neher E. Signal/noise analysis of FRET-based sensors. Biophys J. 2010;99: 2344–2354.

36. Laird DW. Life cycle of connexins in health and disease. Biochem J. 2006;394: 527–543.

37. Butterweck A, Gergs U, Elfgang C, Willecke K, Traub O. Immunochemical characterization of the gap junction protein connexin45 in mouse kidney and transfected human HeLa cells. J Membr Biol. 1994;141: 247–256.

38. Fahrenbach JP, Mejia-Alvarez R, Banach K. The relevance of non-excitable cells for cardiac pacemaker function. J Physiol. 2007;585: 565–578.

39. Cao G, Platisa J, Pieribone VA, Raccuglia D, Kunst M, Nitabach MN. Genetically Targeted Optical Electrophysiology in Intact Neural Circuits. Cell. 2013;154: 904–913.

40. Storace D, Sepehri Rad M, Kang B, Cohen LB, Hughes T, Baker BJ. Toward Better Genetically Encoded Sensors of Membrane Potential. Trends Neurosci. 2016;39: 277–289.

41. Abdelfattah AS, Kawashima T, Singh A, Novak O, Liu H, Shuai Y, et al. Bright and photostable chemigenetic indicators for extended in vivo voltage imaging. Science. 2019;365: 699–704.

42. Rich TC, Tse TE, Rohan JG, Schaack J, Karpen JW. In Vivo Assessment of Local Phosphodiesterase Activity Using Tailored Cyclic Nucleotide–Gated Channels as Camp Sensors. J Gen Physiol. 2001;118: 63–78.

43. Edelstein AD, Tsuchida MA, Amodaj N, Pinkard H, Vale RD, Stuurman N. Advanced methods of microscope control using μManager software. J Biol Methods. 2014;1. doi:10.14440/jbm.2014.36

44. Schindelin J, Arganda-Carreras I, Frise E, Kaynig V, Longair M, Pietzsch T, et al. Fiji: an open-source platform for biological-image analysis. Nat Methods. 2012;9: 676–682.

45. Wu J, Liu L, Matsuda T, Zhao Y, Rebane A, Drobizhev M, et al. Improved orange and red Ca^2^± indicators and photophysical considerations for optogenetic applications. ACS Chem Neurosci. 2013;4: 963–972.

