## Supplementary figures and images for "Optically Activated, Customizable, Excitable Cells - A Kuhl Platform for Evolving Next Gen Biosensors"

### S1 Table

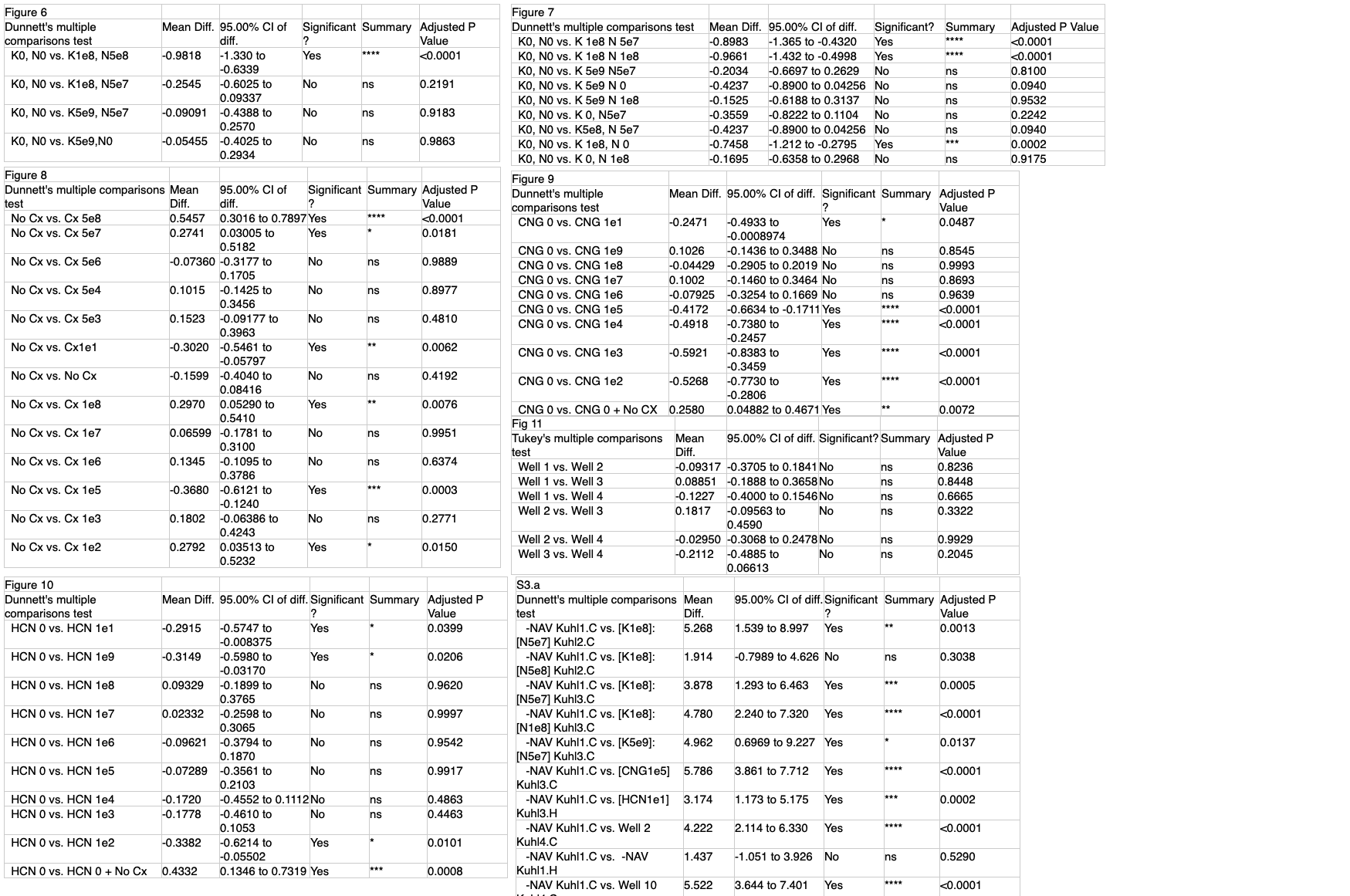

### S2 Table

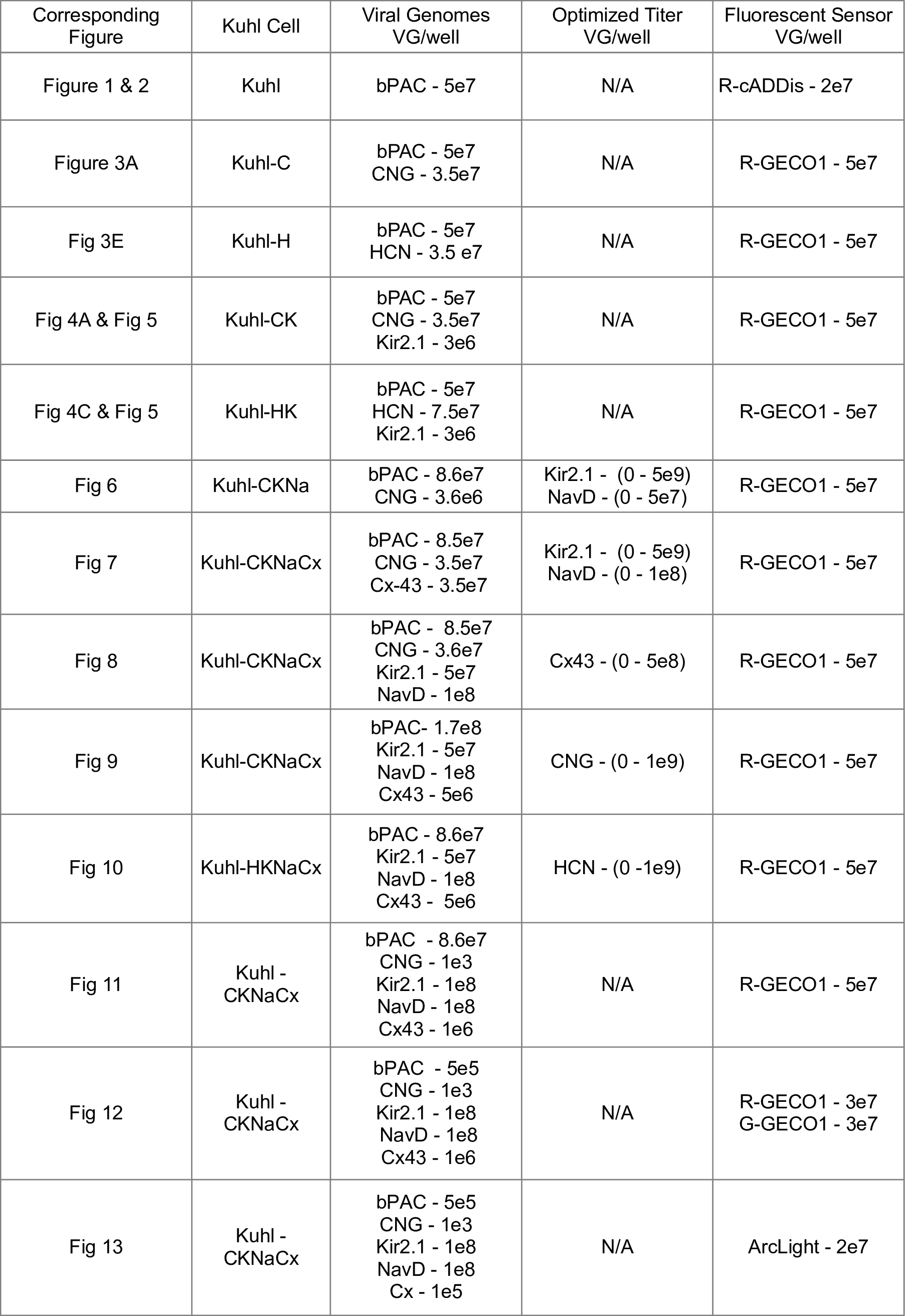

### S3_Fig

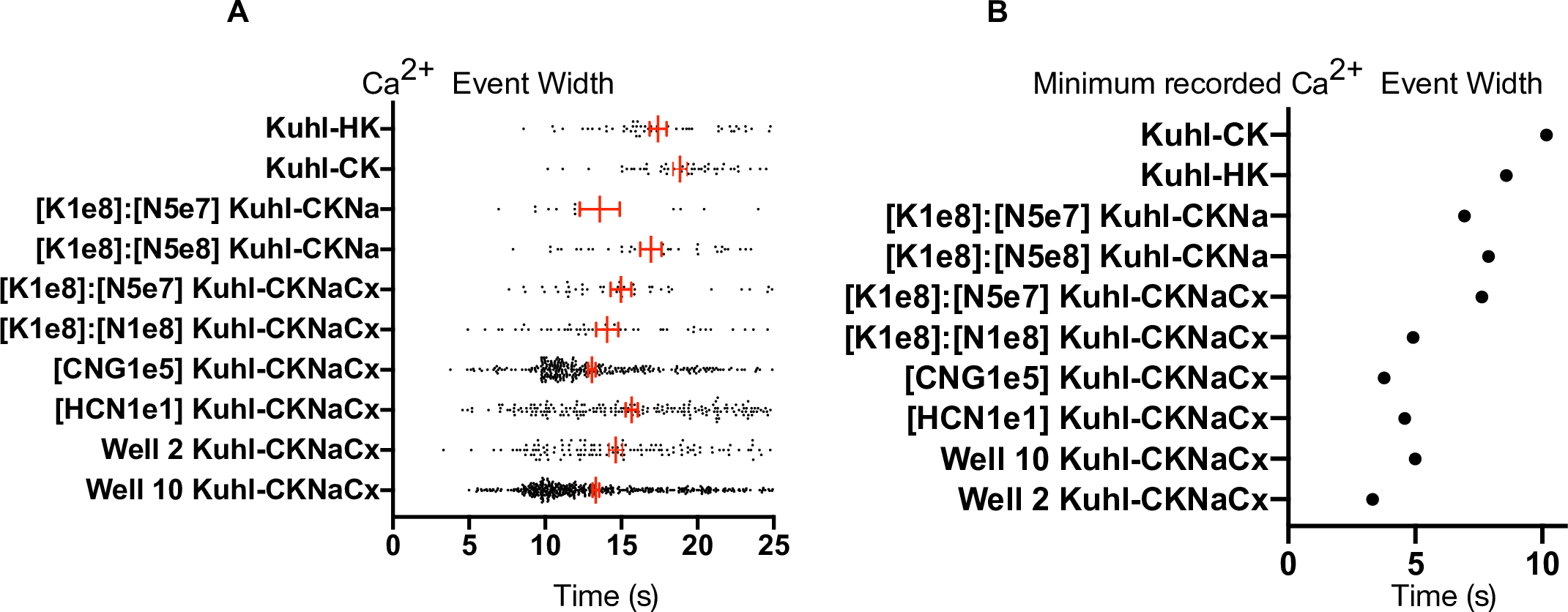

### S4_Fig

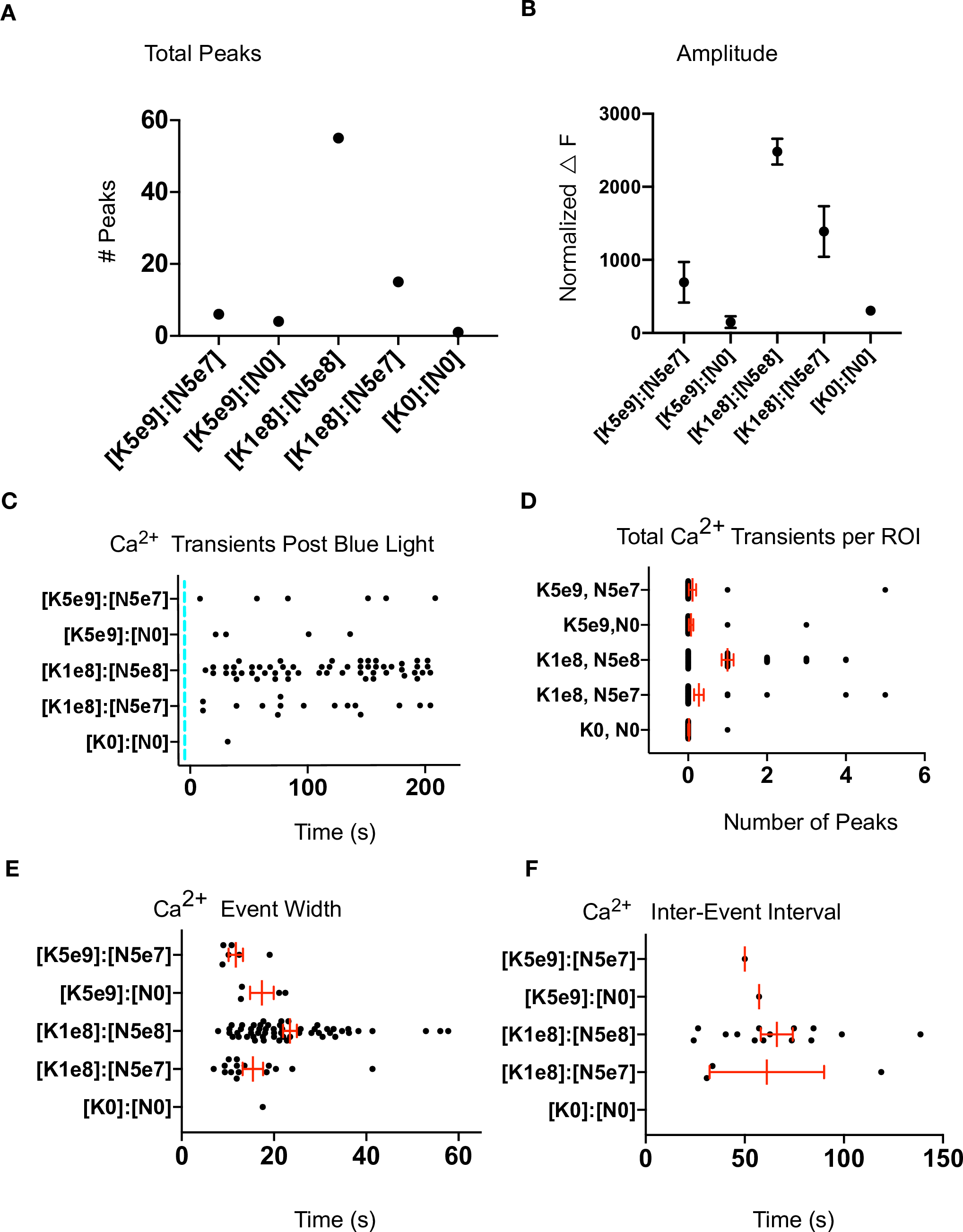

### S5_Fig

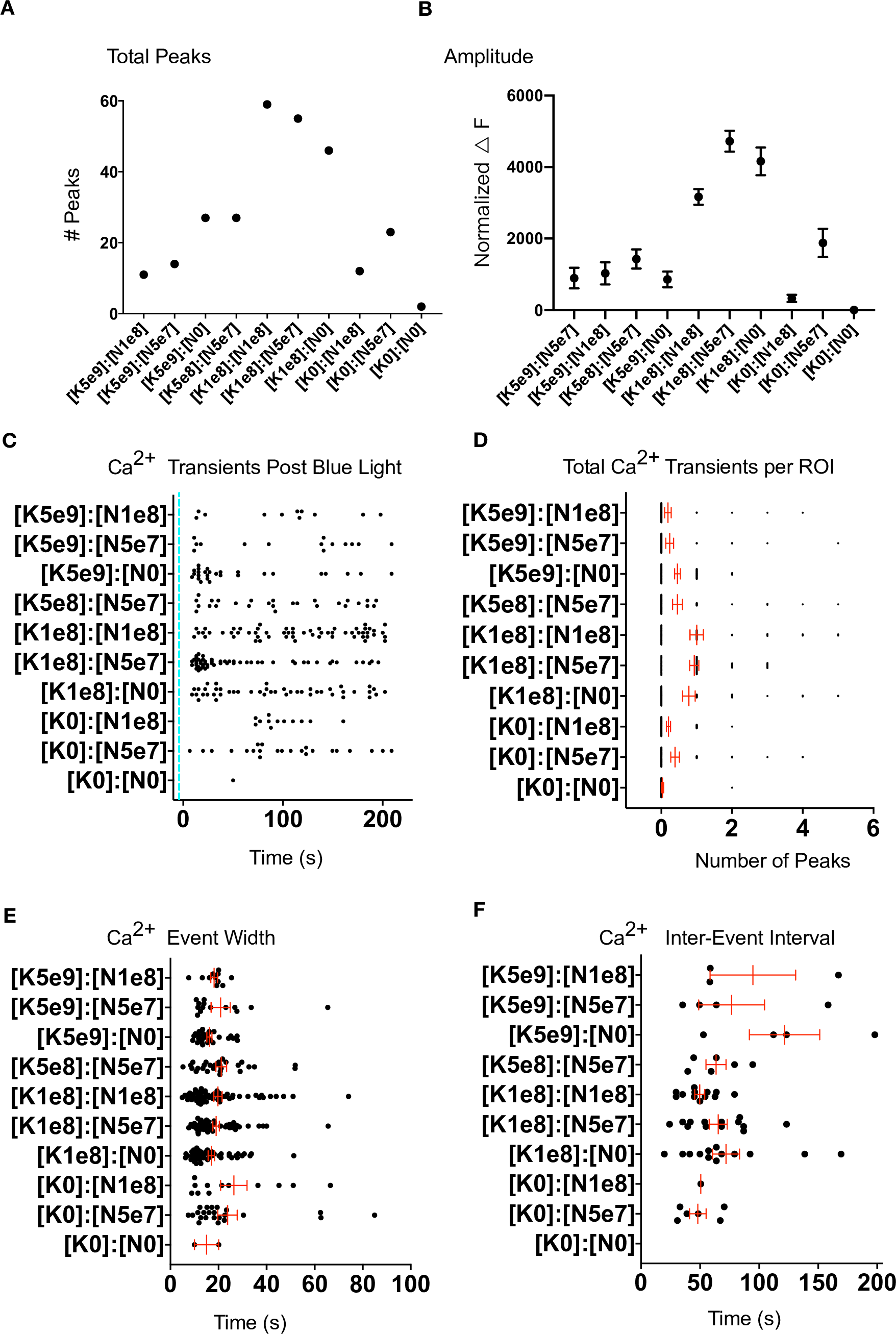

### S6_Fig

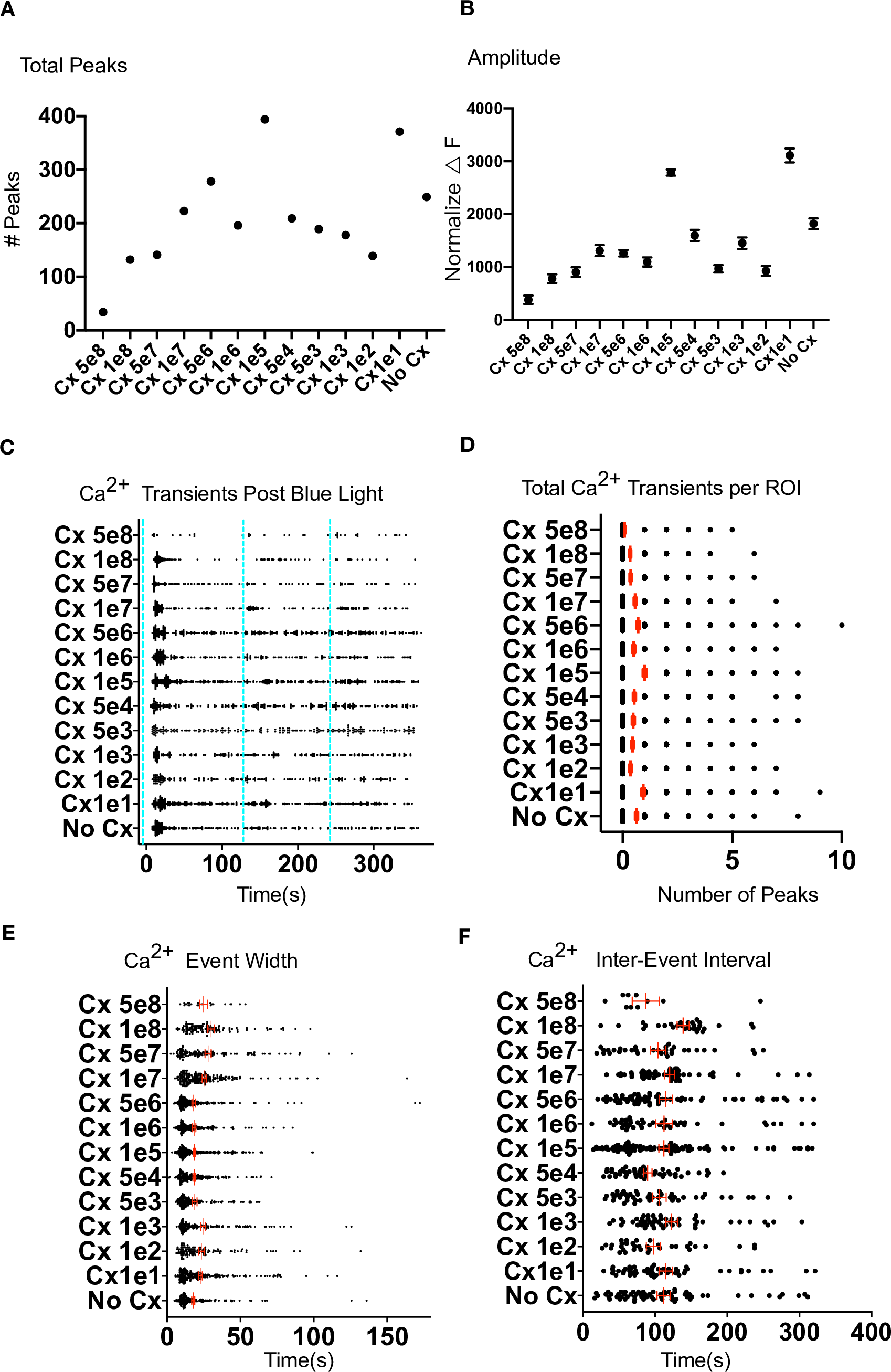

### S7_Fig

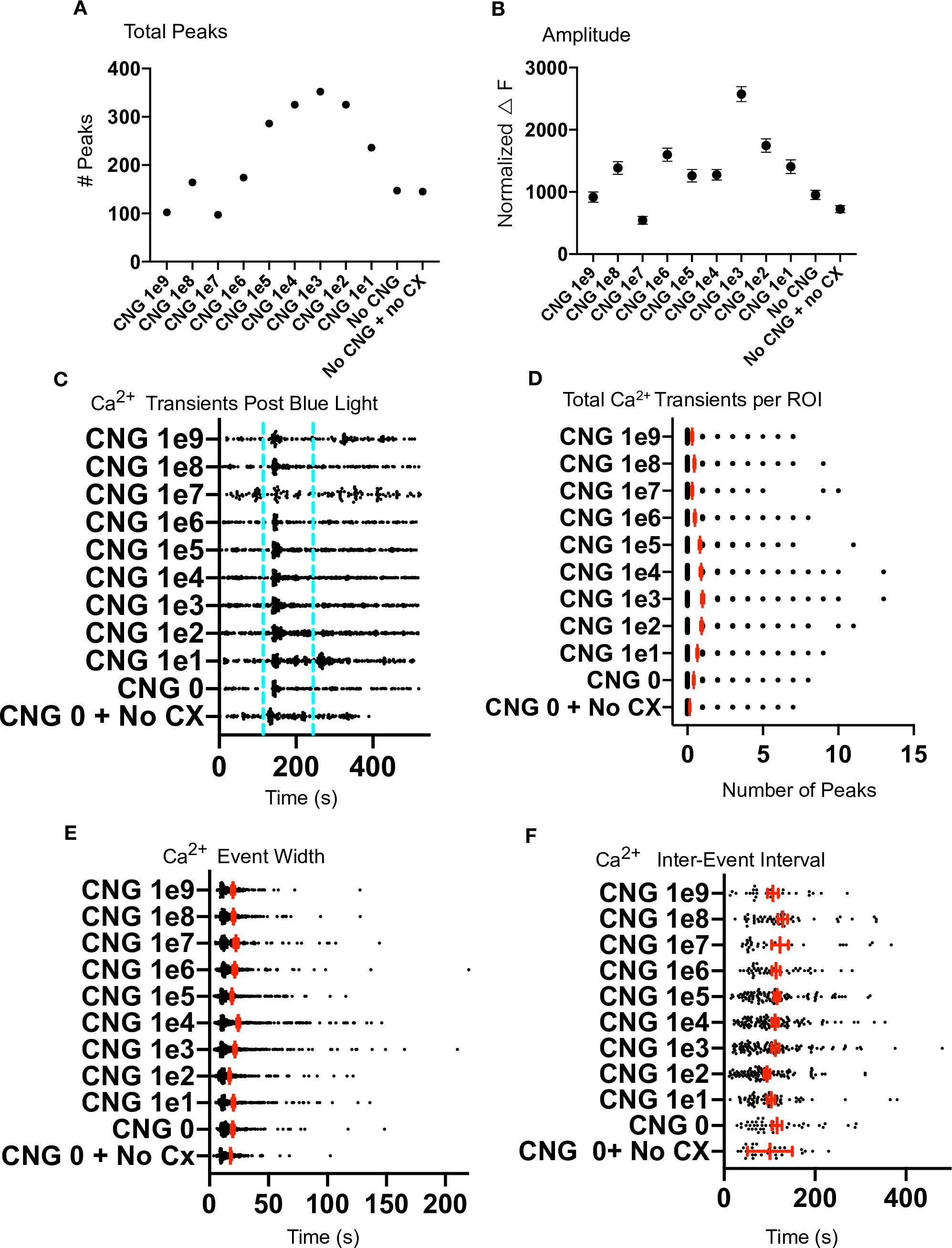

### S8_Fig

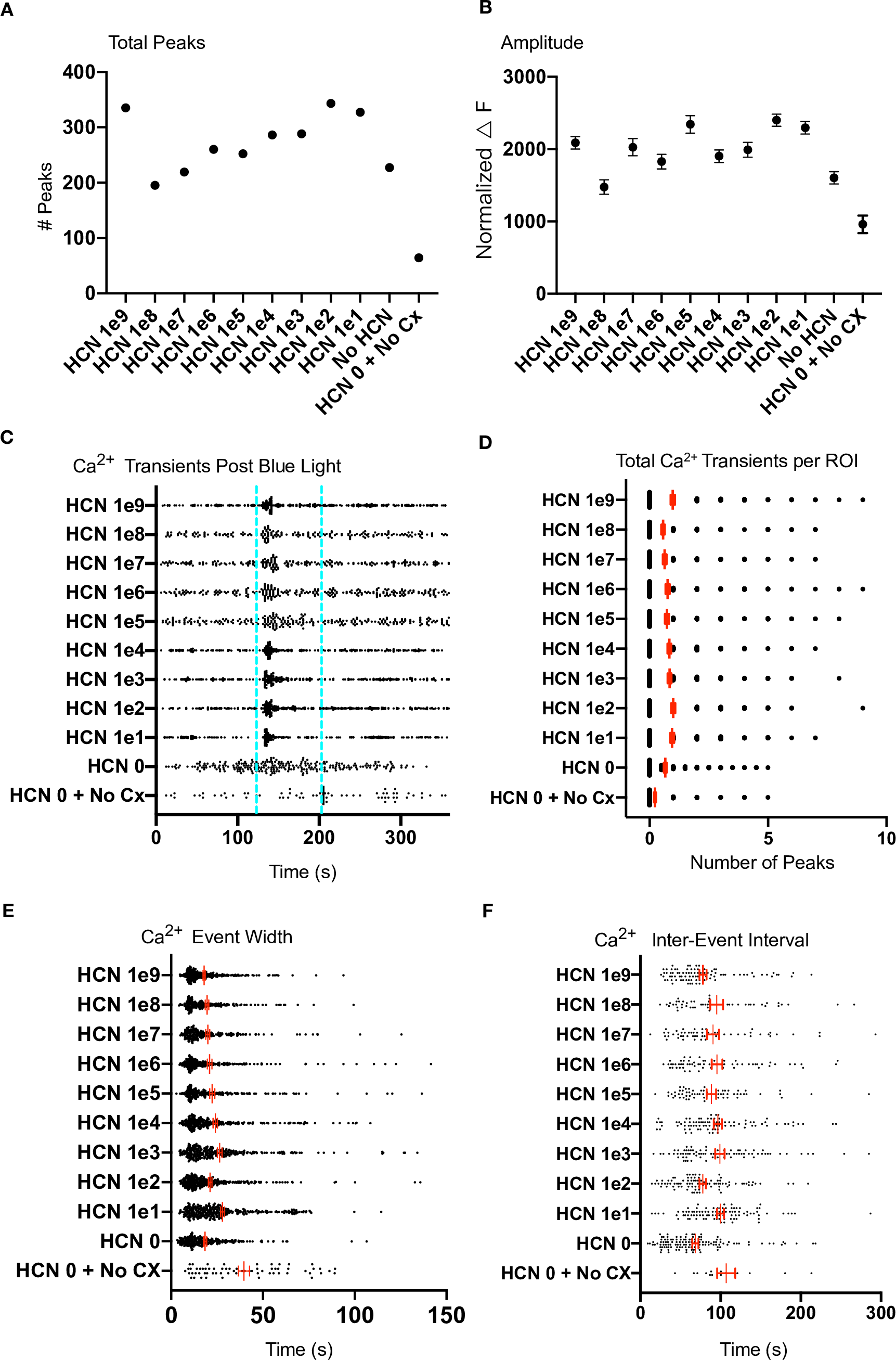

### S9_Fig

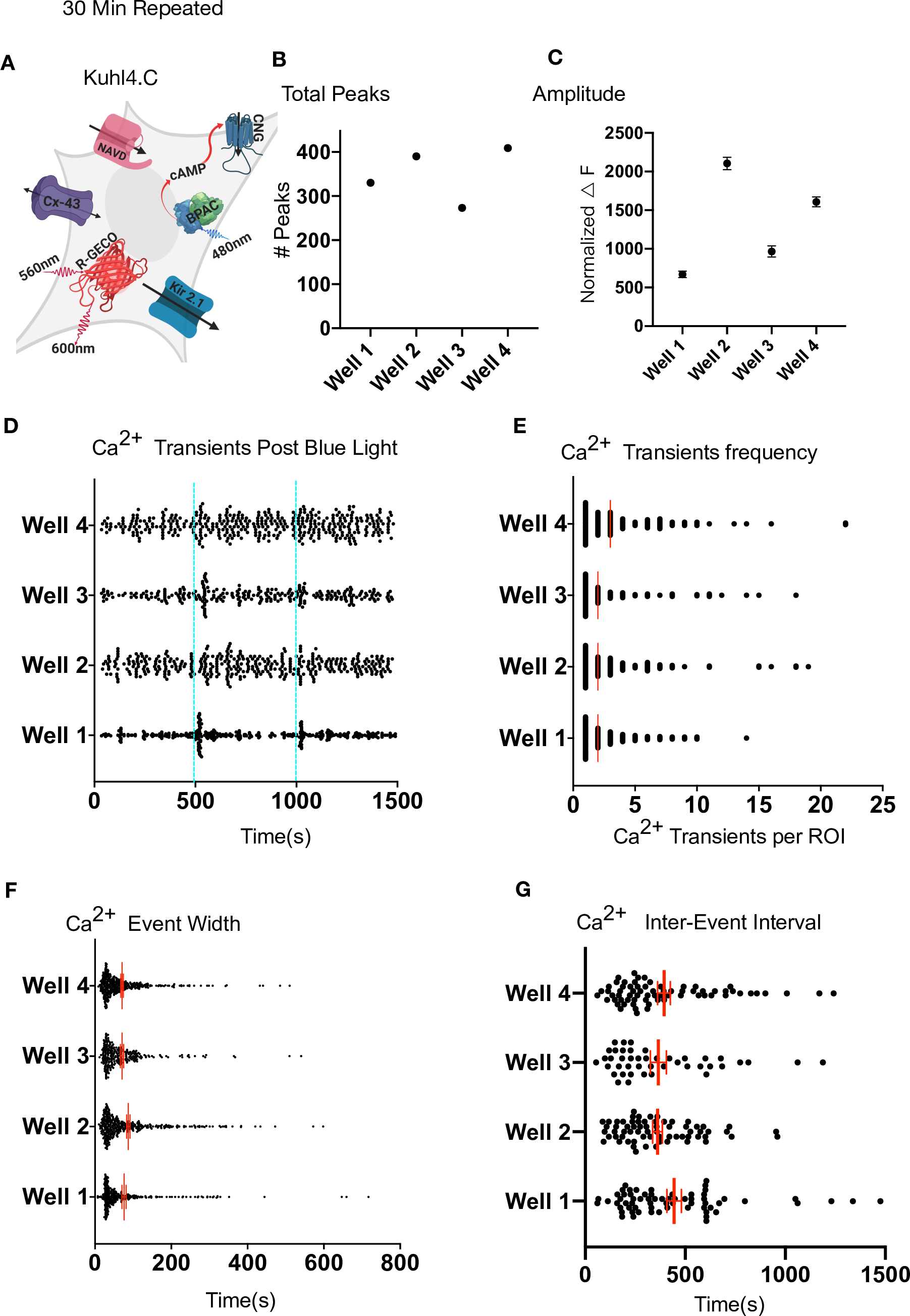

### S10_fig

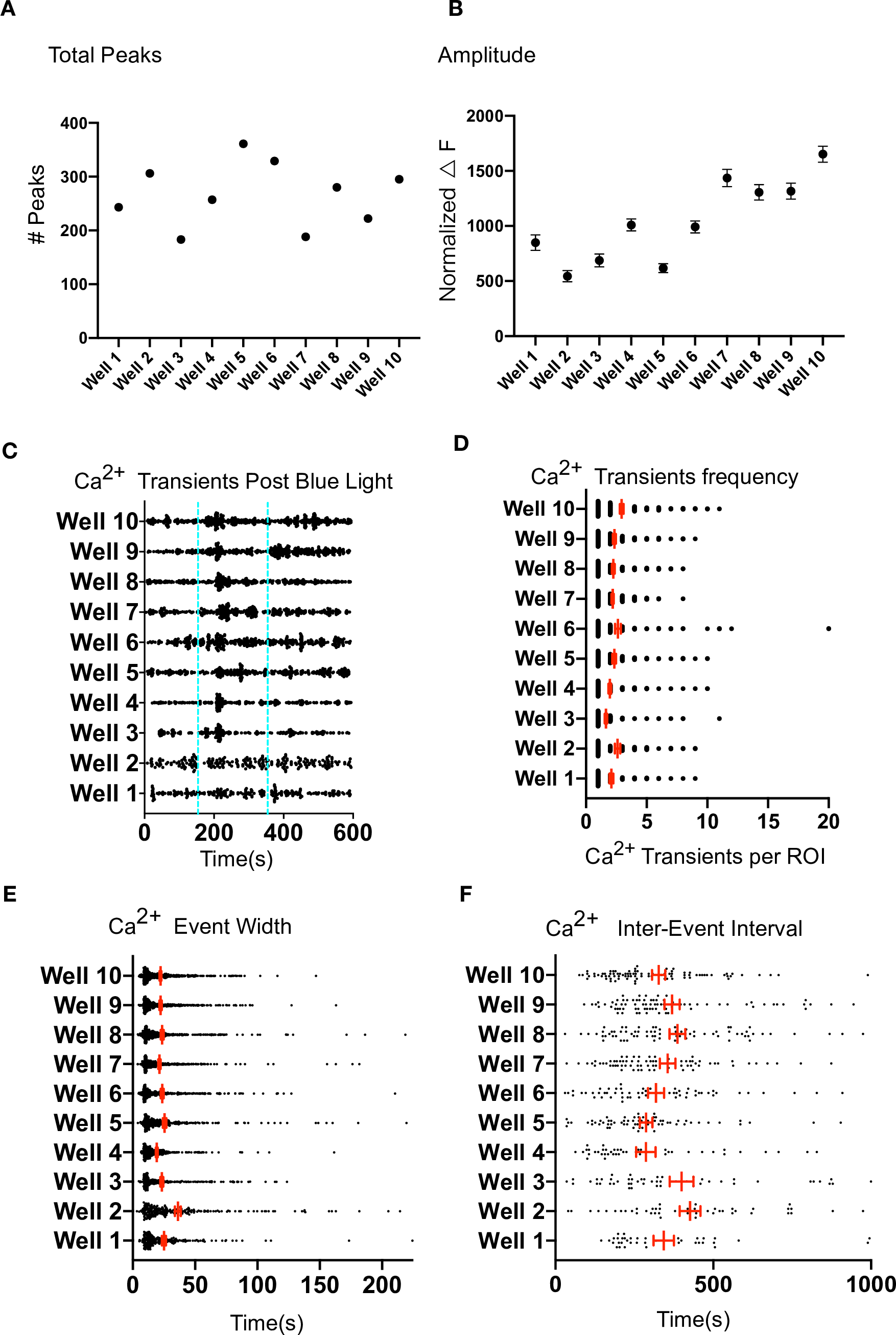

### S Fig 2

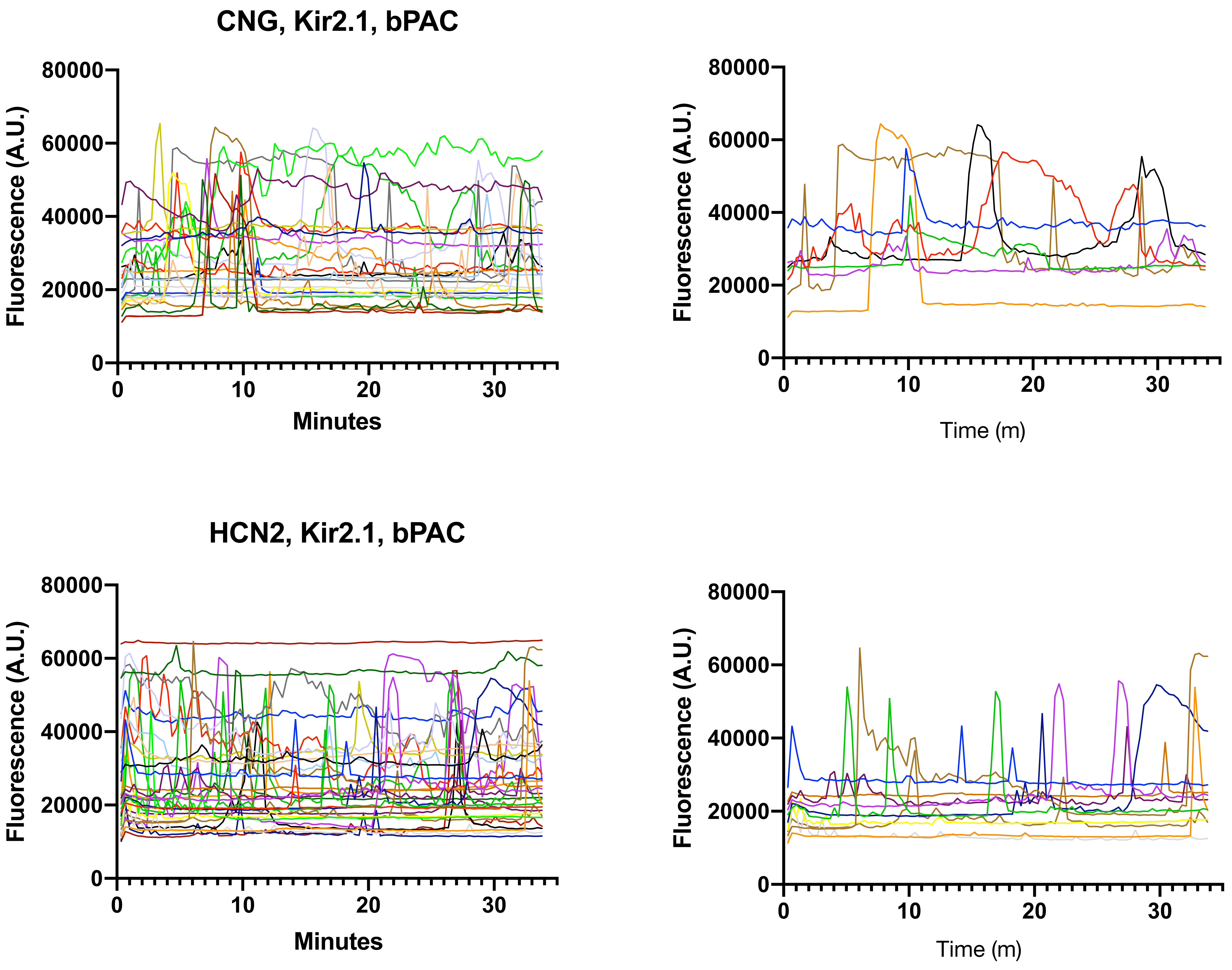

### Supplemental Fig 1

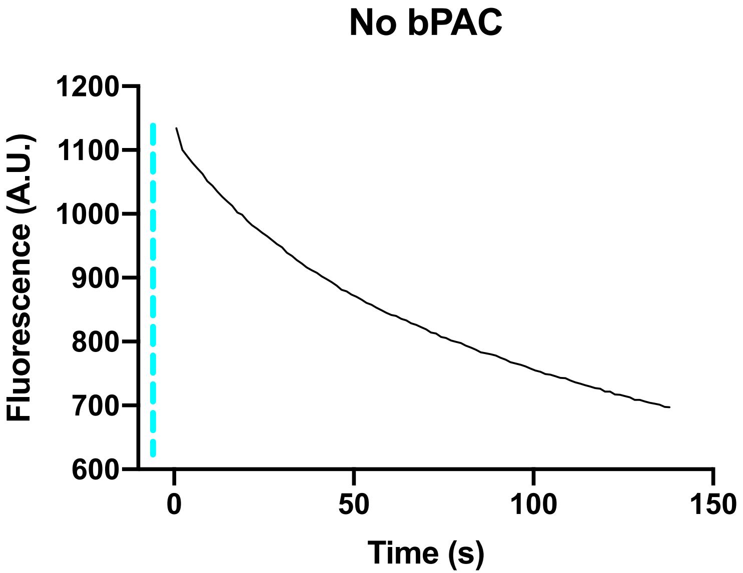

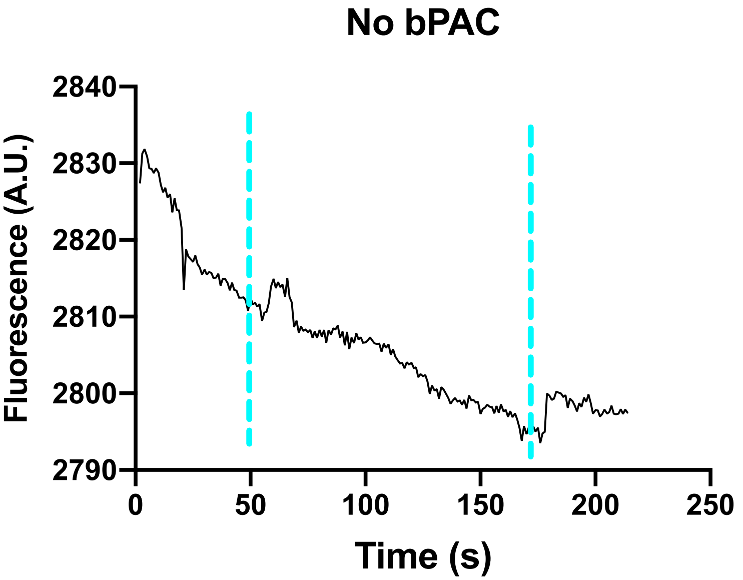
